# Multivariate chemogenomic screening prioritizes new macrofilaricidal leads

**DOI:** 10.1101/2022.07.25.501423

**Authors:** Nicolas J. Wheeler, Kaetlyn T. Ryan, Kendra J. Gallo, Clair R. Henthorn, Spencer S. Ericksen, John D. Chan, Mostafa Zamanian

## Abstract

Development of direct acting macrofilaricides for the treatment of human filariases is hampered by limitations in screening throughput imposed by the parasite life cycle. Efforts to circumvent arduous screening of adult filariae include drug repurposing and high-throughput screens that target commensal bacteria. *In vitro* adult screens typically assess single phenotypes without prior enrichment for chemicals with antifilarial potential. We developed a multivariate screen that identified dozens of compounds with submicromolar macrofilaricidal activity, achieving a hit rate of >50% by leveraging abundantly accessible microfilariae. Adult assays were multiplexed to thoroughly characterize compound activity across relevant parasite fitness traits, including neuromuscular control, fecundity, metabolism, and viability. 17 compounds from a diverse chemogenomic library elicited strong effects on at least one adult trait, with differential potency against microfilariae and adults. Stage-specific drug effects may be crucial to limiting adverse events in endemic regions, and our screen identified five compounds with high potency against adults but low potency or slow-acting microfilaricidal effects, at least one of which acts through a novel mechanism. We show that the use of microfilariae in a primary screen outperforms model nematode developmental assays and virtual screening of protein structures inferred with deep-learning. These data provide new leads for drug development, and the high-content and multiplex assays set a new foundation for antifilarial discovery.

## Introduction

Helminths infect billions of humans, livestock, and companion animals around the world. Vector-borne filarial parasitic nematodes cause two neglected tropical diseases (onchocerciasis and lymphatic filariasis, LF), canine heartworm disease, and various infections of large animals, collectively causing profound devastation across the globe. The portfolio of antifilarials clears the circulating arthropod-infectious microfilariae (mf); however, these drugs do not clear adult worms and some are contraindicated in regions that are co-endemic for multiple filarial nematode species^1^. Concurrent growth of antifilarial use in human medicine, resistance in veterinary medicine^2^, and the paucity of new therapeutic leads present a need for innovation in antifilarial development. Macrofilaricides to treat clinical filariasis cases or sterilizing compounds to interrupt disease transmission are urgently needed^3^.

Screening disease-relevant phenotypes to identify antifilarials is complicated by a number of experimental and biological factors. Screens against adults are particularly encumbered and costly due to their large size, two-host life cycle, and low yield in animal models. In adulticide screening regimens, drug responses are typically measured using single-phenotype *in vitro* assays that are low in resolution and information. Moreover, the extreme phenotypic heterogeneity among infection cohorts and individual *ex vivo* parasites makes assays highly variable^4^. Despite these limitations, there are unrealized opportunities to capture more robust and disease-relevant phenotypic information from individual adult parasites^5^.

An additional opportunity is found in the abundance of mf, which can be isolated in batches of tens of millions from rodent hosts^6^. It is conceivable that a high-throughput primary screen against mf could enrich for compounds with bioactivity against adults. Indeed, >90% of the ∼11,000 genes expressed in adults are also expressed in mf (Supplementary Table 1). Measuring a range of emergent phenotypes in this abundantly accessible pre-larval stage may allow for predictions of general activity against adults. Because chemical perturbation of a common target does not guarantee a conserved phenotypic response in distinct life stages, multivariate screens may help associate disparate drug response phenotypes. For instance, ivermectin diminishes protein secretion by mf and reduces fecundity and motility in adults, the contrasts resulting from stage-specific physiological roles and patterns of localization^7,8^. Multivariate screens with multiple time points could further enhance these strategies by characterizing compounds as “slow” or “fast-acting,” both of which may fulfill different needs in the antifilarial arsenal^9^.

To test these ideas and advance macrofilaricide discovery, we selected a diverse chemogenomic compound library that would allow the exploration of the phenotypic space of mf and adults by targeting classically druggable proteins (Fig. 1a). Each compound in the library is linked to a validated human target, positioning them as molecular probes to discover and validate targets in parasites. In contrast to repurposing screens, the emphasis of a chemogenomic approach is on target discovery in addition to chemical matter. We expect that a subset of compounds with desirable antifilarial properties act through parasite proteins that can be selectively targeted (Fig. 1b). This library was used in a bivariate primary screen against mf, and hit compounds were diverted to a secondary, multivariate screen against adults that parallelized an array of phenotypic endpoints. To inform future anthelmintic screening efforts, we also evaluated the utility of *C. elegans* as a model for antifilarial discovery and the predictive power of virtual screening against human and parasite protein structures modeled by deep learning. We show that tiered, multivariate phenotyping greatly increases the efficiency of hit discovery in macrofilaricide screens and more thoroughly characterizes the bioactivity of lead compounds, resulting in the identification of more than a dozen compounds with submicromolar potency against filarial adults. This strategy, leveraging the unique capacities of distinct parasite stages, could be deployed in other areas of anthelmintic and target discovery.

**Figure 1.**
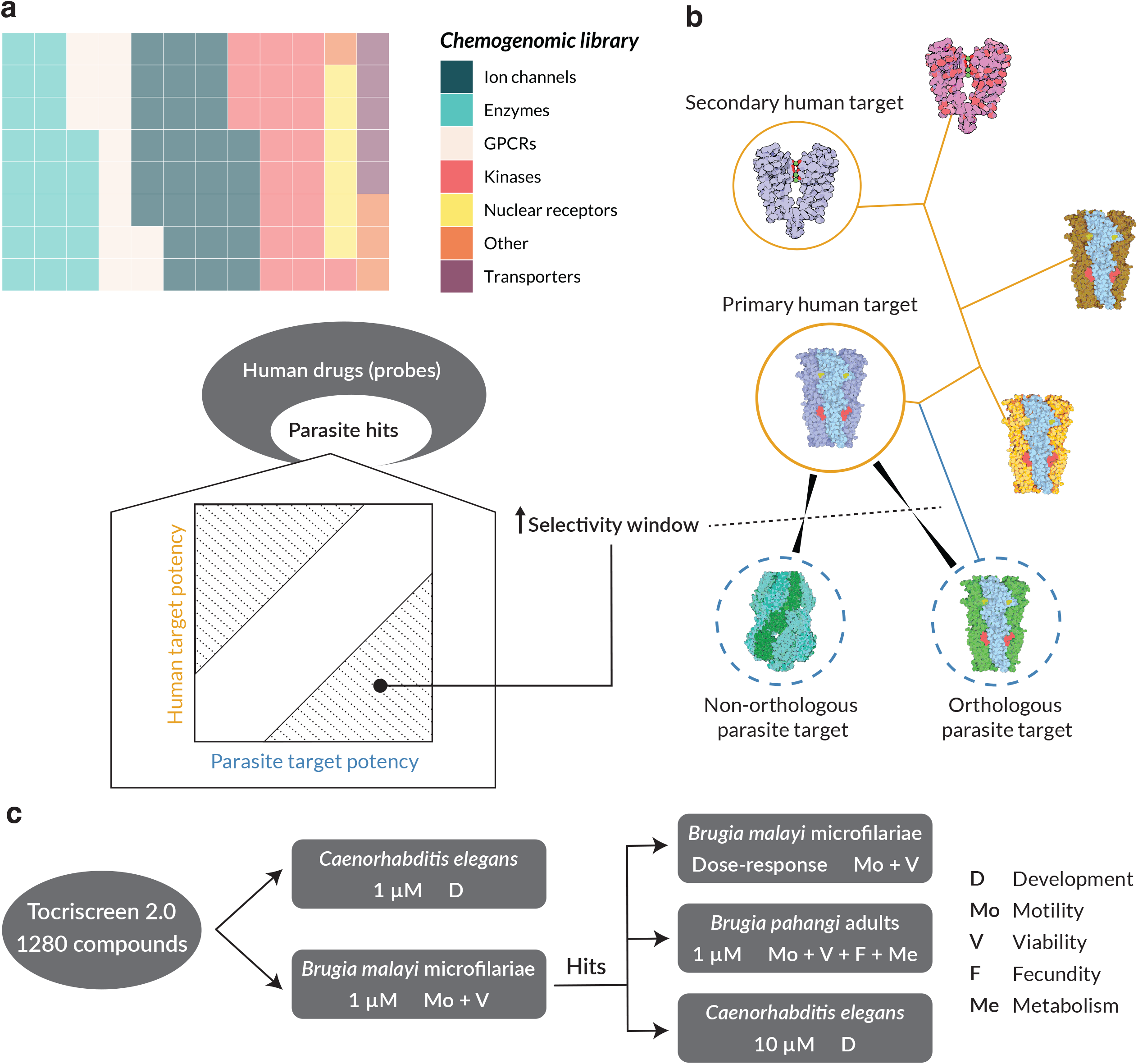
Utilization of chemical probes to identify filarial drug targets. (a) Schematic of how a chemogenomic library can be used to identify non-human drug targets. Chemical probes with known targets are used in a phenotypic screen against parasites. Hits against parasites have a theoretical distribution of relative potencies when compared to the human target; some will be more potent against the parasite target than the human target (bottom right shaded region) and others will be more potent against the human target (top left shaded region). (b) Known phylogenetic relationships among targets may help in deconvoluting the parasite target when it is orthologous to the primary human target, but it may also be the case that the parasite target is specific to the parasite, in which case the known human target will be misleading. (c) Pipeline of the screening approach, incorporating a high-throughput bivariate screen against *Brugia malayi* microfilariae (mf) in parallel to a screen against *Caenorhabditis elegans*. Hits from the mf screen are then followed up for dose-response experiments in mf, a secondary multivariate screen against *Brugia pahangi* adult male and females, and additional screening against *C. elegans* at a higher concentration where necessary.

## Results

### Development of a high-throughput, bivariate microfilariae screen

We developed a bivariate (motility and viability) primary screen that assayed compound activity on microfilariae (mf) at two time points over 36 hours (Fig. 2a-f). We used a high concentration (100 µM) of a diverse compound library to stretch the phenotypic space of mf and allow optimization of a number of different screening parameters, including assay timeline, assay plate design, mf preparation and seeding density, imaging specifications, and image and data processing. Handling and imaging parameters were codified to account for a variety of environmental stimuli that affected mf behavior in plates, including ambient light, room and assay chamber temperature, humidity, settling time prior to imaging, shake speed prior to imaging, and the use of a plate sealer (Supplementary Fig. 1). Deep optimization of imaging-based screens such as this is important for reproducibility across time and space^10^.

**Figure 2.**
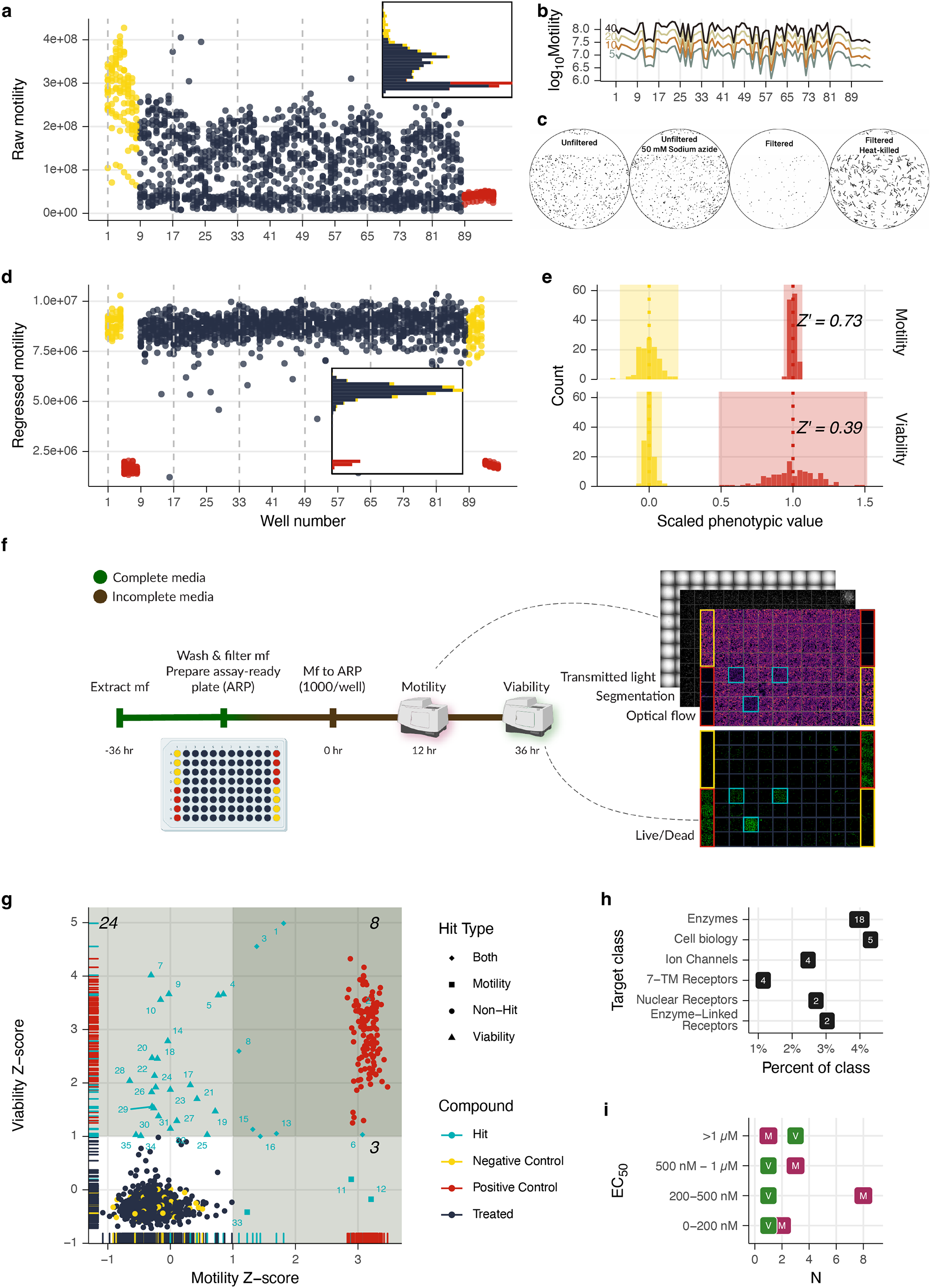
Optimization and employment of a bivariate, high-throughput phenotypic screen of microfilaria. (a) Initial screen of Tocriscreen 2.0 library at 100 µM to optimize plate setup and the number of frames captured. 40 frames resulted in a gradual decrease and patterned oscillation that peaked at the top of each column. (b) Using a reduced number of frames resulted in highly correlated optical flow results. (c) Pre-filtering microfilariae with PD-10 columns and using heat-killing instead of sodium azide as a positive control resulted in a cleaner viability phenotype with reduced embryo and host cell contamination (black pixels are stained material). (d) Pre-filtering mf, regressing raw motility using negative controls at both ends of the plates, and reducing the number of frames corrected the drift in mf movement over the recording period and resulted in overlapping non-hits and negative controls, with greater distance between positive and negative controls (inset). (e) Z’-factors for motility and viability phenotypes (16 plates) after optimization. (f) Final optimized screening strategy. (g) Results of 1 µM screen; Z-score of >1 in each phenotype are shaded. Numbers correlate to enumerated compounds in Table 1. (h) Class annotation of putative hit targets. (i) Binned EC_50_ values for follow-up mf dose-response experiments. M = motility, V = viability.

Parasites from different batches have varying degrees of health and purity, which reduce the signal to noise ratio of the viability assay. We incorporated a column filtration of healthy mf, which substantially cleaned the mf preparations and reduced noise (Fig. 2c)^11^. In our initial optimization experiments, we used sodium azide as a positive control, which had been previously used for viability assays with *C. elegans*^*12*^. However, 50 mM sodium azide paralyzed but did not kill mf, so heat-killed mf were the positive control for all future assays (Fig. 2c).

Initial motility recordings included 40 frames over 15 seconds per well, resulting in a ∼25 minute total acquisition time per plate and a nearly 2x difference in raw motility between the wells at the beginning and end of the plates, caused by mf congregation over time (Fig. 2a). We subsampled data from assay plates and found that the motility calculations from subsampled videos were highly correlated (⍴ > 0.99, Fig. 2b). We chose to acquire 10 frames for each well, reducing the acquisition time and parasite congregation without loss of information. Future data was also normalized using the segmented worm area, and experiments employed a plate design with staggered controls, allowing for regression of values from treatment wells to a line fit across control wells. These acquisition and normalization methods greatly reduced the data variability (Fig. 2d). The final optimized screening strategy included motility measurement 12 hours post-treatment (hpt) and an endpoint viability measurement at 36 hpt. Two phenotypes at alternate time points allowed us to sample drug dynamics, assess potential drug recovery, evaluate phenotypic discorrelation, and potentially reduce the number of false negatives. Z’-factors for motility and viability phenotypes were routinely greater than 0.7 and 0.35, respectively (Fig. 2e)^13^.

### Identification and characterization of hit and target combinations with nanomolar potency against microfilariae

We used this optimized bivariate strategy to screen the Tocriscreen 2.0 library at 1 µM against *B. malayi* mf (Fig. 2f). The library contains 1,280 bioactive compounds with pharmacological classes that have been theorized as anthelmintic targets, including GPCRs, kinases and other enzymes, ion channels, and nuclear receptors, among others^14–16^. The diversity of targets and their broad physiological roles increased the likelihood of obtaining hits associated with diverse phenotypes. While pursuing hit compounds with known activity in humans may enable drug repurposing, we are also interested in leveraging this information for the validation of parasite targets that could be diverted into mechanism-based screening strategies (Fig. 1a-b).

The primary screen identified 35 hits (Z-score > 1; 2.7% hit rate) (Fig. 2g). While the overall correlation between the two phenotypes was high (*r =* -0.84), there was less correlation among our hits (*r =* 0.33), reflecting the capture of non-redundant phenotypic information. Indeed, only 32 (91%) or 11 (31%) hits would have been identified using only viability or motility alone, respectively. We presume that the later time point (36 hpt vs 12 hpt) of the viability assay is the primary reason for the greater number of hits, but also note that of the 11 motility hits, only eight were viability hits. Although these phenotypes often reinforce each other, they can also be decoupled by the action of particular drugs. The 35 hits span a range of target classes (Fig. 1h, Table 1). Interestingly, four of the hits are histone demethylase inhibitors, and two act on the NF-κB/IκB pathway. Out of the human targets, 16 contain homologues in *B. malayi*, 12 of them one-to-one, and the human targets of primary hits were not more likely to have more similar parasite homologs than the targets of non-hits (Supplementary Data 1, Supplementary Fig. 2).

We generated eight-point dose-response curves for 31 of these hits, recording motility at 12 hpt and both phenotypes at 36 hpt. 15 hits had a motility or viability phenotype at 36 hpt (>25% reduction in viability or motility when compared to control, Supplementary Fig. 3a). As expected, the liberal threshold set in the primary screen resulted in some compounds not reproducing an effect when retested. Six of eight compounds that produced effects in both motility and viability phenotypes in the primary screen reproduced effects in the dose-response experiment, while 2/3 and 7/19 of the motility and viability hits were reproduced, respectively. Thirteen of the hits had EC_50_ values of <1 µM for at least one phenotype, and ten of these were <500 nM (Fig. 2i). Two hits had an EC_50_ of less than 100 nM: NSC 319726, a p53 reactivator (19 nM) and NAV 2729, an Arf6 inhibitor (81 nM). These two compounds caused substantial motility and viability effects (Z_Viability_ > 4.5, Z_Motility_ > 1.3), suggesting that our primary screen had some predictive power for compound potency. Eleven of the top 15 hits from the primary screen were recapitulated with <1 µM EC_50_ values, while only 3 of the bottom 22 hits produced a measurable EC_50_ for any of the phenotype/time point combinations. Indeed, there was a significant correlation between the EC_50_ for motility at 36 hpt and Z_Viability_ + Z_Motility_ (*r* = -0.594, p-value = 0.0249, df = 12).

Dose-response experiments with measurements at multiple time points allowed us to distinguish between “fast-acting” compounds (Phenotype_12hpt_ == Phenotype_36hpt_) and “slow-acting” compounds (Phenotype_12hpt_ < Phenotype_36hpt_). We do not assume that a good anthelmintic lead compound (or target) should induce a fast effect. Slow deterioration of dead worms, leading to a slower release of antigens and commensal *Wolbachia* bacteria, may serve to reduce adverse effects after drug treatment^9,17^. Six compounds had effects at 12 hpt that were similar to the effects at 36 hpt (area between the curves < 0.3) while the remainder had a higher effect at 36 hpt (Supplementary Fig. 3b). Two of the seven fast-acting compounds are NF-κB/IκB inhibitors.

### Virtual screening has a modest ability to enrich for bioactive compounds and predicts orthologous targets with dissimilar binding pockets

Utilization of computational filters or surrogate screening models prior to a primary screen, with the goal of increasing throughput and efficiency, is a consistent theme in antifilarial research and development. With data from our primary screen in hand, we investigated whether these alternative approaches could have identified the same hit compounds as those resulting from our primary screen, have allowed us to screen fewer compounds, or help direct towards the likely parasite targets of our hits.

We first tested the ability of virtual screening to recall known interactions. We docked each compound in the Tocriscreen library to a set of 3,284 AlphaFold2 (AF2) models of human proteins (4,194,311 total docking runs), which included the known target(s) of each compound. If this strategy could successfully extract known interactions, we could use it to generate leads for the parasite targets of hit compounds. We used AF2 models and blind docking in anticipation of translating the approach to the parasite, for which AF2 models exist but have unannotated binding sites. We tuned machine learning models to classify actives and decoys based on the known interactions with human targets. A tuned random forest achieved a 6-fold enrichment factor at 1% when evaluated on hold-out test data (that is, we were 6 times more likely than random to identify an active in the top 1% of probabilities generated by the classifier), a robust initial enhancement (RIE^18^) of 5.29, and a BEDROC^19^ of 0.14. These metrics indicate that our virtual screening strategy has a modest ability to recall compound-target interactions.

In addition, we tested whether this computational strategy could classify the hits that were identified in our bivariate phenotypic screen. We repeated the blind docking strategy with 828 parasite target structures, orthologs of the human targets used in the previous analysis, and used the tuned model to predict actives/decoys and predict the target for each hit compound (Supplementary Table 2). Our virtual screening strategy would have slightly enriched for actives, but any pre-filtration would have also led to a substantial dropout of hits (Fig. 3a). For instance, screening the 640 compounds that had the highest maximum docking probability (the best 50%) would have recovered 18 (53%) of the primary hits (Fig. 3a).

**Figure 3.**
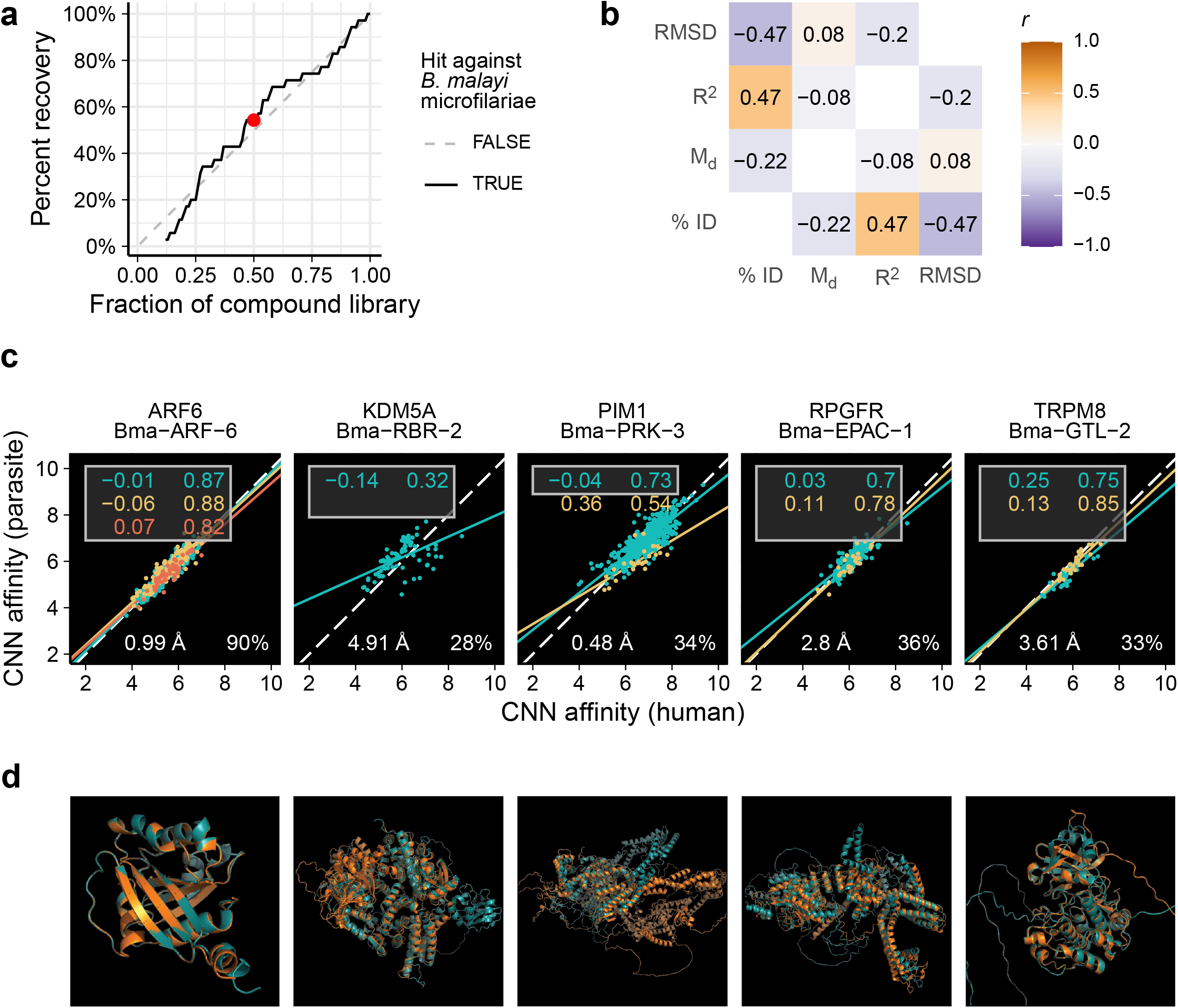
A virtual screening strategy using AlphaFold2 models of *B. malayi* and human targets. (a) Performance of a virtual screening strategy as a filter prior to an *in vitro* primary screen. When compounds are ranked by docking score, using only the highest ranked compounds at different cutoffs would have slightly enriched for hits against microfilaria (e.g. screening the highest ranked 50% of the library would have resulted in retrieval of 60% of the hits - red point). (b) A docking-based binding site comparison between orthologous host and parasite targets provides unique information that is non-redundant with other types of comparisons. (c) Clockwise starting at the top-left: mean distance from y=x (M_d_), R^2^ of linear fits, primary sequence percent identity, and root-mean-square deviation of atomic position of aligned AlphaFold2 structures. Lines, points, and labels are colored by binding site (only the top 3 consensus sites were retained, site #1 = cyan, site #2= yellow, site #3 = red). Each point represent the affinity scores for the same compound docked to orthologous structures. (d) Aligned AlphaFold2 orthologous structures from *B. malayi* (cyan) and humans (orange).

Finally, the virtual screens allowed for comparison of the docking profiles of human and parasite orthologs as an assessment of binding pocket dissimilarities. Several approaches have been used to assert the existence of a potential selectivity window when investigating anthelmintic protein targets with known host orthologs, including primary sequence similarity and structural alignment^20–22^. However, ortholog pairs can have low sequence identity while having highly conserved binding pockets, and structural alignment followed by assessment of binding pocket dissimilarity requires *a priori* target selection and knowledge of the binding site. We designed a metric that measured binding pocket structural dissimilarity between host/parasite orthologs using high-throughput blind docking (Fig. 3b). This metric (M_d_) summarizes the deviation of site-specific docking affinities between orthologous pairs from the assumption that they would be identical for identical binding pockets. M_d_ poorly correlates with other metrics of dissimilarity such as the root-mean-square deviation of atoms in aligned orthologous structures and the percent identity of aligned orthologous primary sequences, demonstrating that this metric provides unique information (Fig. 3b). Among putative targets of our primary screen hits, KDM5A/Bma−RBR−2 had the highest M_d_, while ARF6/Bma−ARF−6 had the lowest (Fig. 3c-d). This metric leverages advances in deep learning for protein structural determination and molecular docking to generate genome-wide predictions of the ability to selectively target the parasite proteins where host orthologs can be identified.

### *C. elegans* development poorly predicts antifilarial activity

The model nematode *C. elegans* has been used as a surrogate for anthelmintic screening efforts with mixed results, especially when used as a model for nematodes that reside in separate phylogenetic clades^23,24^. Nonetheless, *C. elegans* is a convenient screening model to discover nematicidal compounds and offers genetic tools that have enabled the discovery of the targets of such compounds^25–29^. To evaluate *C. elegans* as a primary screening model for antifilarial discovery, we assayed the effects of the Tocriscreen library on *C. elegans* development. Four compounds caused a development reduction one standard deviation away from the mean (0.3% hit rate), and only one hit (NAV 2729) was shared with the hits from the *B. malayi* mf primary screen (Fig. 4a). For this library, a *C. elegans* primary screening filter would have led to the discovery of only one of 35 compounds that have activity against *B. malayi* mf.

**Figure 4.**
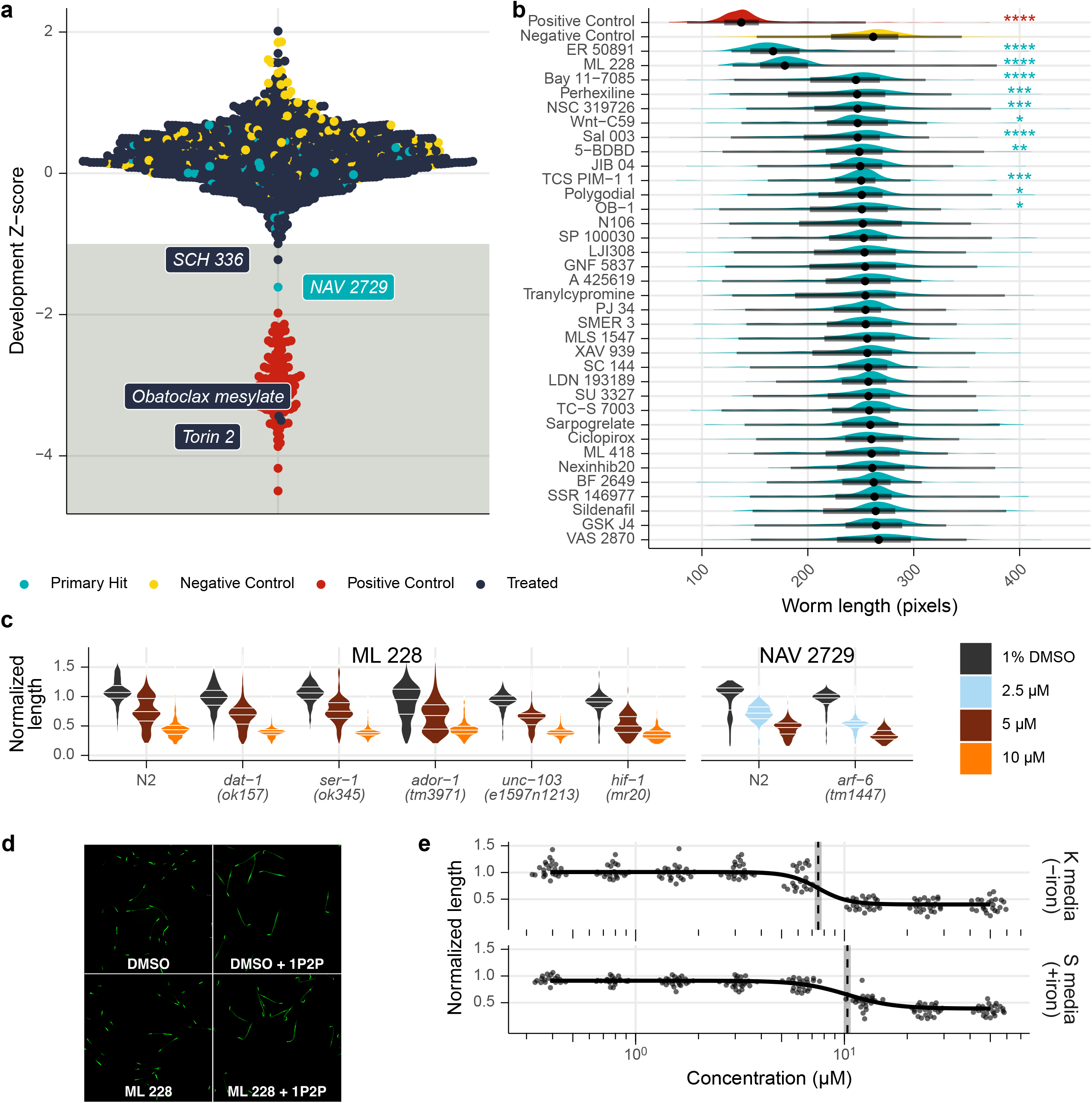
A development screen of a free-living model nematode. (a) *C. elegans* development screen of entire Tocriscreen 2.0 compound library at 1 µM. Light blue hits are shared in *B. malayi* mf (see Figure 2). (b) Follow-up 10 µM screen of hits in *B. malayi* mf that were not hits in *C. elegans* at 1 µM. Positive control is 50 mM albendazole sulfoxide. Asterisks indicate significance (****: p < 0.0001, *** = p < 0.001, ** = p < 0.01, * = p < 0.05; Dunn’s test). (c) *C. elegans* development assay with selected knock-out strains. Data are normalized to the median length of the DMSO-treated worms. (d) Hypoxia-activation after treatment with 50 µM ML 228 or 1.25% 1P2P. (e) ML 228 dose-response of *C. elegans* development when grown in K or S media.

*C. elegans* has a thick collagen cuticle that is impermeable to polar and large molecules, and high concentrations of compound are often necessary to enable pharmacologically relevant bioaccumulation^30,31^. To test the possibility that the thicker cuticle of *C. elegans* masked the discovery of antifilarial compounds, we exposed *C. elegans* to a higher concentration (10 µM) of the subset of compounds that were active against *B. malayi* mf. An additional 11 compounds caused significant developmental defects at 10 µM, but only two caused effect sizes >30% (ER 50891 and ML 228, Fig. 4b). These data clearly show that a *C. elegans* development screen provided little predictive power for identifying compounds with *in vitro* bioactivity against filaria, even when exposing worms to comparatively higher concentrations and longer exposure times.

*C. elegans* has been utilized as a platform for deconvoluting the target or mechanism of action of a hit shared between free-living and parasitic nematodes^29^. We attempted to identify the target of compounds that exhibited activity against both *C. elegans* and *Brugia* mf. We chose NAV 2729, the Arf (GTPase) inhibitor that had potent effects on mf motility and viability and ML 228, a hypoxia-inducible factor (HIF) activator. We repeated *C. elegans* development assays with NAV 2729 or ML 228 treatment and included strains that had knock-outs for putative targets^32,33^. We hypothesized that absence of the predicted drug target, which we selected by using the phylogenies of the known human target (Supplementary Fig. 4), would result in the abrogation of the drug-induced developmental delays.

In the presence of drug, each knock-out strain experienced the same developmental delay as wild type, suggesting that these targets are not the primary mediators of the drug effect observed in *C. elegans* (Fig. 4c). ML 228 is believed to induce a hypoxia response in human cell lines by chelating iron, causing HIF-1ɑ activation^32^. The developmental effect of ML 228 on *C. elegans* persisted even in the *hif-1* knockout strain, suggesting that ML 228 does not activate a hypoxia response in nematodes. Consistent with this hypothesis, 50 µM ML 228 did not cause increased GFP expression in a hypoxia reporter (Fig. 4d). Since ML 228 does not exert its developmental effect via HIF activation, we wondered if the effect was due to a generalized iron chelation. In our developmental assay, worms acquire iron from their bacterial food source. We performed dose-response experiments in S medium, which has slightly different salt and cholesterol contents and is supplemented with trace metals including iron. Growth in S medium resulted in a 38% increase in EC_50_, which may be attributed to the extra iron (Fig. 4e). In human cell lines, iron supplementation resulted in a 13-fold increase in ML 228 EC_50_^32^.

These data show that genetic knockout of orthologous candidate targets did not modulate drug susceptibility in *C. elegans*. It is therefore likely that both NAV 2729 and ML 228 exert their antinematodal activity through non-orthologous targets associated with different mechanisms of action to what is known in mammals.

### Optimization of a multiplex adult screening regimen

*Brugia pahangi*, a parasite of felids that is closely related to the etiological agents of LF, can be reared in rodent hosts at higher yields than *B. malayi* and thus has been used as a surrogate in motility-based screens^34^. To optimize a multilex phenotyping strategy for macrofilaricides, we compared the activities of control and heat-killed *B. malayi* and *B. pahangi* during *in vitro* culture over a period of 5 days. Individual worms remain highly motile over this time period and produce measurable amounts of lactate (Fig. 5a-b). Female fecundity of both species showed a monotonic decrease over time, with *B. pahangi* being significantly more fecund than *B. malayi*, making it a more sensitive organism for measuring drug-induced inhibition (Fig. 5c-d). Females that started with low fecundity never increased in fecundity, allowing the formulation of a convenient filter for pre-treatment health. Most females stopped laying mf after 48 hrs. in culture, and heat-killed worms were incapable of laying progeny (Fig. 5e). These experiments also tested the effects of serum on worm health. Serum can have non-random effects on pharmacodynamics and can interfere with assays on conditioned media, making incomplete (i.e., serum-free) media the preference. Serum supplementation had no effect on motility over 5 days (Fig. 5a), but it did slow the decrease in fecundity (Fig. 5c).

**Figure 5.**
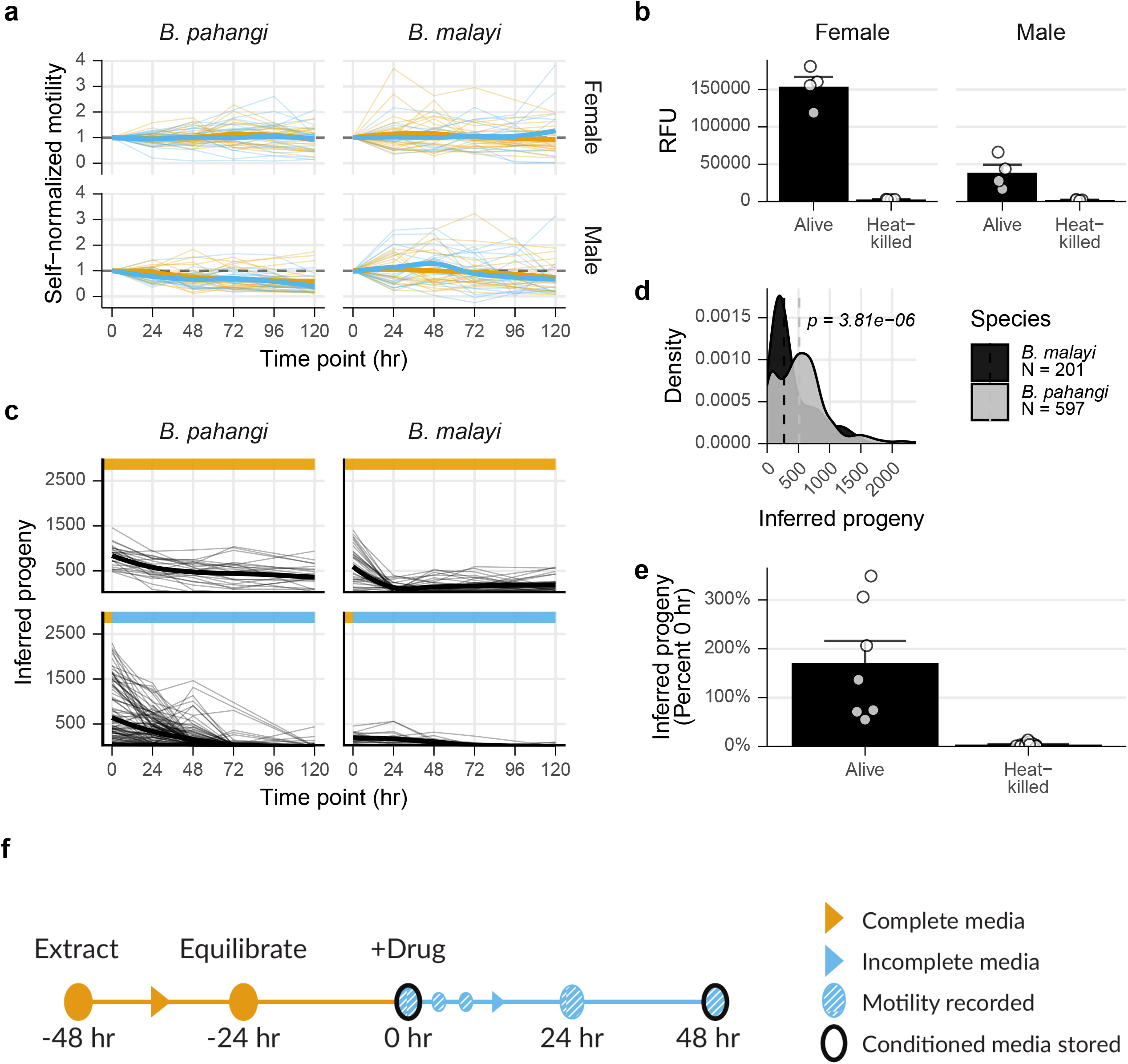
Optimization of multivariate *Brugia* spp. adult screen. (a) Motility of *B. pahangi* and *B. malayi* adult males and females over 120 hours in culture in media with (complete, orange) or without (incomplete, blue) serum. (b) Lactate production of *B. pahangi* adult males and females after 48 hours in incomplete media. (c) Progeny released every 24 hours over 120 hours in culture in complete or incomplete media. (d) Progeny released after overnight culture in complete media. P-value calculated with Kruskal-Wallis test. (e) Progeny released after 48 hours. (f) Final optimized screening strategy incorporating four total phenotypes.

*Ex vivo* filarial adults exhibit substantial phenotypic heterogeneity and batch variability. Female batches can vary in fecundity by three orders of magnitude (Supplementary Fig. 5), and both sexes display a complex variety of body postures and knotting behaviors. In order to control for this variation, we ensured that phenotypes could be self-normalized to a pretreatment value after overnight equilibration in complete media, followed by a 48 hr assay timeline (Fig. 5f).

### Identification of hit and target combinations with sub-micromolar activity against adult filarial parasites

Thirty-one of the hits from the primary screen were tested at 1 µM in this optimized secondary screen. Motility was measured before and immediately after drug addition, and at 1, 24, and 48 hpt. Fecundity was measured before drug addition and at the conclusion of the experiment. Metabolic activity was assessed via measurement of lactate produced in the conditioned media, and worms were terminally stained with CellTox to measure potential adulticidal activity of the compounds.

We defined a motility hit as any compound that causes >50% decrease in motility in over half of the tested worms, or an average decrease of 75%; similar cutoffs were used for the other phenotypes (see Methods). This heuristic ensured that a single worm that showed a stochastic increase in motility did not cause false negatives when the other worms all exhibited substantial decreases in motility. Thirteen compounds satisfied this threshold after 48 hours, a hit rate of 32% (Fig. 6a-b). Four of these compounds caused motility reductions in both sexes, 3 in females only, and 6 in males only. The motility hits against females only accounted for 6 of the 11 compounds that were most effective in causing a reduction in fecundity (Fig. 6c). That is, 5 compounds had a fecundity effect on females but did not have a motility effect. Eight compounds caused reductions in lactate release by females, and 4 caused reductions in males. In females, all the motility hits were also metabolism hits, two hits were only metabolism and fecundity, and one was a metabolism only hit. In males two hits were metabolism hits but not motility hits, seven hits were motility only hits, and two were hits in both phenotypes (Fig. 7a). In total, 17 compounds (55%) elicited a hit in at least one phenotypic assay, with female fecundity being the most sensitive phenotype (11 hits). These data clearly show that the additional multiplexed phenotypes reveal information about compounds that would not have been captured using motility alone.

**Figure 6.**
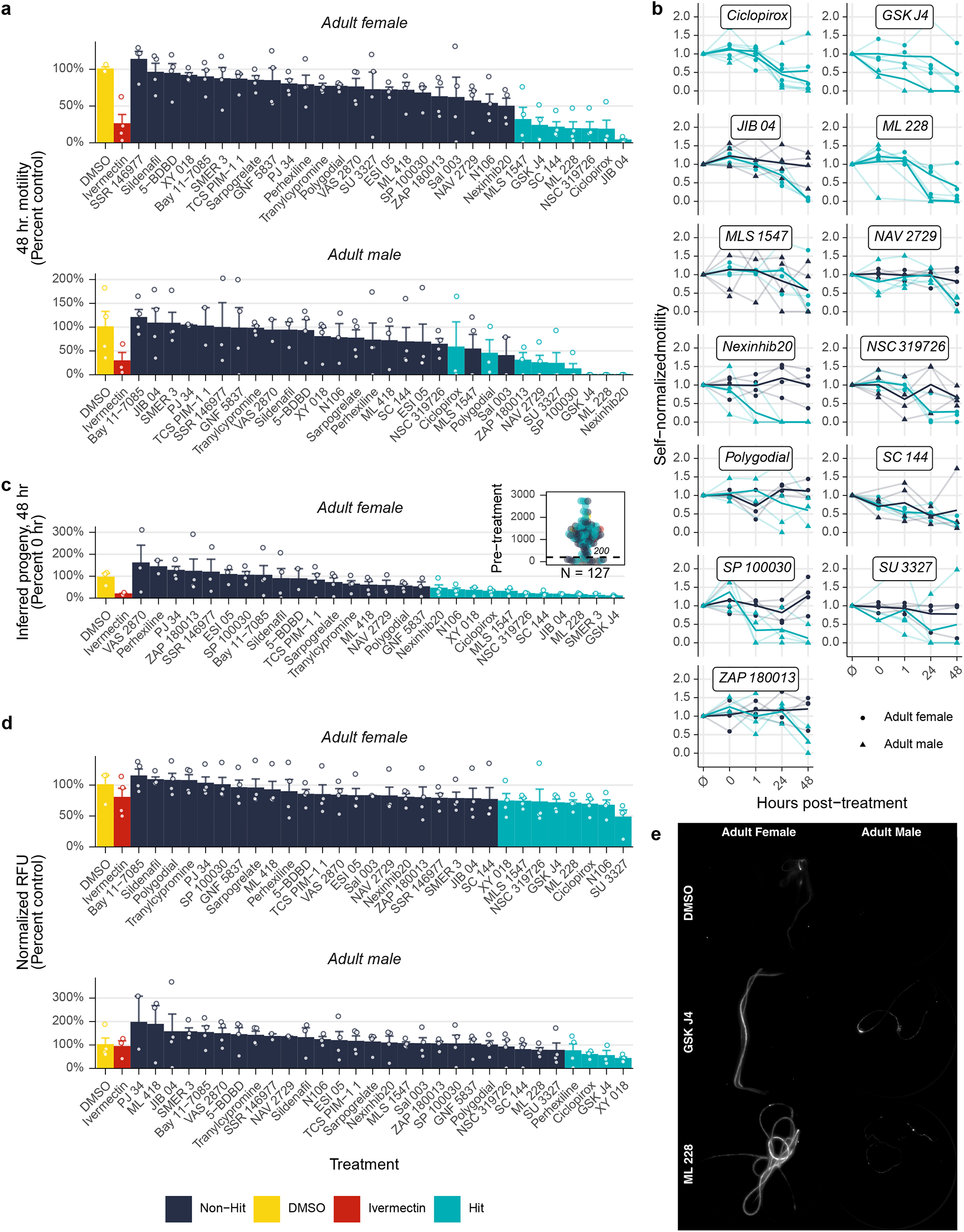
Secondary screen of adult male and female *B. pahangi*. (a) Motility of worms after 48 hours post-treatment (hpt). Compounds in light blue are hits, navy blue are not hits. (b) Individual worm motility over time. Ø = immediately prior to drug addition, 0 = immediately after drug addition. (c) Progeny released 48 hpt. (Inset) Female worms that released less than 200 mf after overnight culture in complete media were filtered from downstream analysis. (d) Lactate production after 48 hpt. (e) Example images of viability-stained adults. Stained worms are dead. For all bar charts, each point is an individual worm, columns are the mean of the group, and bars represent the standard error of the mean.

**Figure 7.**
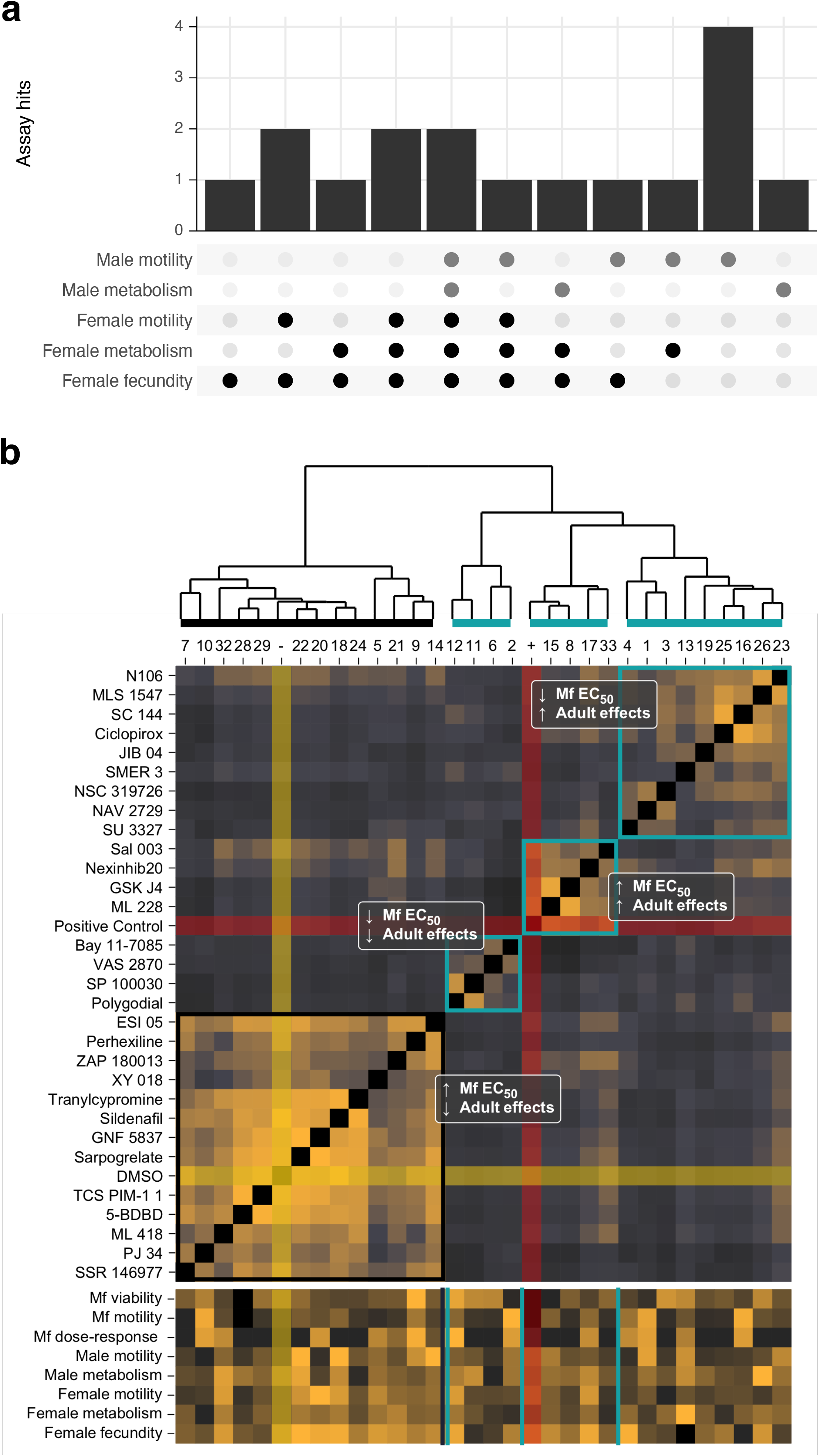
Antifilarial hits cluster into four phenotypic groups. (a) Tally of the number of hits from each assay combination. For instance, two compounds were hits in all five assays, and four compounds were hits in the male motility assay alone. (b) Hits against microfilaria (mf) in the primary screen but failed to reproduce in the followup dose-response and had no effects against adults (black rectangle) clustered with the negative control (yellow highlight). Four compounds were potent against mf but had no effect on adults, five compounds were relatively less potent against mf but had strong effects on adults, and the remaining hits (nine compounds) were potent against mf and adults (blue boxes).

Two compounds (Ciclopirox and GSK J4) had effects across all five phenotypes, and ML 228 had an effect on four phenotypes (Fig. 7a). Ciclopirox and GSK J4 are histone demethylase inhibitors (KDMIs), and a third KDMI (JIB 04) had a female-specific effect. KDMIs have a rich history as antischistosomal leads^35–37^, and have been recently identified in macrofilaricidal screening regimens^38^. Our data add to this corpus of evidence, and, interestingly, our two-stage screening approach suggests that these epigenetic drugs may be potentially lucrative as specific adulticidal targets. All four of the KDMIs that were identified in the primary screen poorly reproduced in dose-response experiments, adding to the evidence that although they have effects against mf, they are much more potent against adults^38^. Furthermore, GSK J4 and ML 228 are likely adulticidal, as some treated worms were brightly stained by a cytotoxicity stain (Fig. 6e).

We collated the data from all the parasite screens and clustered the hits based on scaled phenotypic values (Fig. 7). Hits were classified into four broad categories: 1) those with high EC_50_ values on mf and no effects on adults (14 compounds), which clustered with DMSO; 2) those with high EC_50_ values on mf and potent effects on adults (5 compounds, macrofilaricidal), which clustered with ivermectin; 3) those with low EC_50_ values on mf and no effects on adults (4 compounds, microfilaricidal); 4) those that were potent against both mf and adults (9 compounds). A summary of all screening data can be found in Table 1. The most interesting compounds characterized by this thorough set of analyses include the aforementioned ARF6 inhibitor, which is lethal to mf has an EC_50_ on motility of 81 nM, decreases *C. elegans* by more than 50% at 5 µM, and reduced male motility by 75% at only 1 µM; the four histone demethylase inhibitors, which have parasite-specific effects and greatly reduce female fecundity; the two NF-kB/IkB inhibitors, which have highly potent effects on motility but reduced effects on adults, and ML 228, which is more potent on adults than mf and caused significant effects on several adult phenotypes. However, one of the unique strengths of our approach is the ability to illuminate diverse bioactivities of different hit compounds, allowing investigators to prioritize compounds associated with diverse mechanisms linked to different emergent phenotypes of relevance to *in vivo* efficacy.

## Discussion

The development of macrofilaricides is a key component to eliminating filarial Neglected Tropical Diseases and was recently recognized as a critical action in the World Health Organizations 2021-2030 roadmap^39^. The movement to utilize *in vitro* screening of parasites as the primary driver for macrofilaricide development has yielded few leads even with massive investment. Targeting the endosymbiont *Wolbachia* to sterilize and slowly kill adult worms is an alternative strategy, and doxycycline has been successfully used in a test-and-treat approach for onchocerciasis^40^. High-throughput screening has recently identified a number of anti-*Wolbachia* leads that exhibit desirable pre-clinical traits, some of which are now in clinical trials^41–44^. However, compounds that act directly on parasites are still highly desirable, even if leveraging high-throughput technologies to identify them are hindered. Repurposing anthelmintics from veterinary medicine for the treatment of filariases is an additional ongoing effort, and emodepside is currently advancing through clinical trials for the treatment of onchocerciasis, but is not viable as a treatment for LF^45^.

We developed and deployed a novel strategy to identify direct-acting macrofilacides, incorporating a highly optimized primary screen against microfilaria to enrich for compounds with antifilarial activity. Primary hits were then characterized for activity against adults by inspecting an expanded repertoire of multiplexed disease-relevant phenotypes, in contrast to traditional single-endpoint screens. This tiered approach greatly reduced the time and investment required for hit identification and substantially increased the amount of information gleaned about a compound of interest. Even though the primary screen was carried against mf, it resulted in the identification of compounds with greater potency against adults than mf, which is a desirable property in regions where rapid mf death may result in severe adverse events^1^. Likewise, utilizing multiple timepoints in the primary and secondary screens allowed us to prioritize drugs based on the preference for slow or fast-acting compounds and/or compounds that cause permanent or transient, and therefore potentially recoverable, effects.

Chemogenomic approaches in which compounds are used as molecular probes linked to known host targets, provide leads for parasite target identification and have been used to great effect in other infectious disease systems, including *Plasmodium falciparum* and *Mycobacterium tuberculosis*^46,47^. *However, in contrast to these systems, the genetic tools used to discover and validate targets are underdeveloped in filarial parasites. Use of the genetically tractable model nematode C. elegans* allowed us to rule out the predicted targets of two drugs with conserved activity across filarial and free-living nematodes. Although we had anticipated mapping drug effects to host-orthologous targets, our results reveal a more desirable trait: compounds with potent antifilarial activity that are most likely mediated by completely novel mechanisms. For example, NAV 2729 was highly potent against mf and had strong male-specific effects on adults; but these actions are not mediated by the nematode homolog of the known human target (ARF6). The same is true of ML 228, a compound that had strong effects against both adult sexes within 24 hours and was comparatively less potent against mf, but for which the HIF homolog is not the nematode target.

For the antifilarial compounds that also cause developmental delay in *C. elegans*, forward genetic screens can be utilized to identify the nematode target instead of a candidate gene approach. Mapping the targets of compounds with specific antiparasitic action, such as the multiple mammalian KDM and NF-κB pathway inhibitors, will require a different approach. Reverse genetics approaches have been used to validate anthelmintic targets^22,48,49^, as have candidate gene approaches based on the known interactions between the compound and host targets^50^. These could be similarly leveraged to elucidate targets of our hit compounds, though an RNAi-based strategy would only be available for adult hits and would be highly target-dependent.

Virtual screening of structural models inferred by deep-learning had limited utility in target identification, but it provided a method of assessing structural dissimilarity between host and parasite orthologs. Off-target host toxicity is an ever present concern when dealing with multicellular eukaryotic parasites, and outside of traditional medicinal chemistry and comprehensive structure-activity relationship studies, which require heterologous expression of the target and a relevant high-throughput screening assay, few approaches have been developed to assess genome-wide structural dissimilarity between host-parasite orthologs. Methods based on primary sequence similarity have proliferated in tandem with the assembly of new helminth genomes^51^, but the pharmacological relevance of these approaches remains unclear. In contrast, our approach using high-throughput molecular docking generates affinity profiles for each potential binding pocket and allows for the unbiased, genome-wide determination of targets with the most dissimilar docking profiles. This approach provides unique information that can be incorporated when evaluating and selecting drug targets from genomic data sources.

We present here a comprehensive optimization of phenotypic and computational screening strategies for antifilarial drug and target discovery. We identified compounds with stage, species, and/or sex-specific activity, some with submicrolar potency. The phenotypic spectrum of each hit compound is thoroughly profiled through the use of parallelized multivariate assays, providing a rich dataset that allows the selection of compounds with distinct traits rather than characterizing a compound simply as an “active.” Each compound is linked to a putative target, and the use of model organisms and virtual screening of structural models provides leads for target identification. These data seed future compound and target-centric investigation of diverse hit profiles, and the methods developed provide an advanced strategy for filarial phenotypic screens and antifilarial discovery.

## Methods

### Parasite maintenance

*Brugia malayi* and *Brugia pahangi* life cycle stages were obtained through the NIH/NIAID Filariasis Research Reagent Resource Center (FR3); morphological voucher specimens are stored at the Harold W. Manter Museum at University of Nebraska, accession numbers P2021-2032^52^. Parasites were maintained at 37°C with 5% atmospheric CO_2_ in RPMI 1640 culture media (Sigma-Aldrich, St. Louis, MO) with penicillin/streptomycin (0.1 mg/mL, Gibco, Gaithersburg, MD) and FBS (10% v/v, Gibco) unless otherwise stated.

### Screening of *Brugia malayi* microfilariae

A chemogenomic compound library of 1,280 bioactive small molecules (Tocriscreen 2.0, Tocris Bioscience, Minneapolis, MN) was stored at 1 mM in DMSO in low dead volume 384-well plates. Assay-ready plates were generated at the UW-Madison Small Molecule Screening Facility with a Labcyte Echo 650 (Beckman Coulter, Brea, CA) liquid handler by spotting each compound onto 96-well assay plates (Greiner Bio-One, Frickenhausen, Germany), which were sealed with adhesive foil, wrapped in parafilm, and stored at -20°C. The entire library was screened against *B. malayi* microfilariae (mf) at 100 and 1 μM after a single freeze-thaw cycle. A detailed procedure for performing the primary mf screen is reported at: [Protocol Exchange link].

Videos were analyzed with the motility and segmentation modules of wrmXpress v1.0^53^. Videos were analyzed on a node maintained at UW-Madison’s Center for High Throughput Computing (CHTC). Output data was analyzed using the R statistical software, including tidyverse packages and drc to analyze dose-response experiments^54,55^. Motility data was processed by normalizing flow values to the log_2_ of the segmented worm area, corrected using a linear model that included well row and column as numeric predictors, and scaling between 0 and 1 using the means for the negative (1% DMSO) and positive (heat-killed) controls, respectively. Outliers (identified using the IQR method with a coefficient of 3) in positive and negative controls were removed. Hits were defined as any treatment that had a Z > 1.

### Screening of *C. elegans*

*C. elegans* N2 (Bristol) were maintained at 20°C on NGM plates seeded with *E. coli* OP50 and routinely picked to fresh plates at the L4 stage. Gravid worms were bleached and embryos were hatched in K media (or S media for iron chelation experiments) overnight (14-20 hours) at a concentration of 3 embryos/μL. Hatched L1s were titered by counting 10 aliquots of 5 µL spots, and K or S media was added to adjust the suspension to 1 L1/μL. Food solution was made by diluting frozen *E. coli* HB101 stock (50 mg/mL) 10X in K or S supplemented with kanamycin (Fisher Scientific AC450810100, Waltham, MA).

For the 1 µM library screen, assay-ready plates were prepared with 1000X compound, and the first/last column of every plate contained final concentrations of 1% DMSO or 50 µM albendazole sulfoxide (Fisher Scientific). 50 μL of the titered L1 culture was added to each well followed by 50 μL of food solution. Plates were sealed with breathable film and incubated at 20°C in a humid chamber with shaking at 180 RPM. At 48 hpt, each well was filled with sodium azide (final concentration 50 mM, Fisher Scientific) or 1-phenoxy-2-propanol (1P2P, Sigma-Aldrich, final concentration 0.7%) to paralyze the worms, and entire wells were imaged at 2X with an ImageXpress Nano (Molecular Devices).

Images were analyzed with the worm size module of wrmXpress^53^. Segmented worms were computationally straightened, and the length of each worm was used to measure development.

### Multivariate screening of adult *Brugia pahangi*

A detailed procedure for performing the multivariate adult screen is reported at: [Protocol Exchange link]. Care was taken to work quickly and keep parasites at physiological temperature, as we have observed substantial differences in worm behavior when handled at room temperature^56^. Each compound was tested in quadruplicate against both sexes, and plates were recorded in a custom imaging setup prior to adding drug, immediately after adding drug, and 1, 24 and 48 hpt (Fig. 3f). At 48 hpt, worms were picked from 24-well plates to 96-well plates with 100 µL of PBS per well and stained for viability.

### Adult *Brugia pahangi* motility analysis

Videos of individual wells were cropped and motility was measured with a previously described optical flow algorithm^57^. For optimization experiments, worms with 0 hr. flow values of less than 5% of the minimum value or less than the background flow were annotated as dead and removed from any further analysis, as were any flow value that was >2.5x that worm’s 0 hr. flow value. For screening experiments, the raw optical flow was log_10_ transformed, and the background flow was subtracted using the mean flow of empty wells. Videos were also manually inspected and nonmoving worms at the 0 hr. time point were removed from further analysis. A hit was defined as any compound that caused a decrease of >50% in motility in over half of the tested worms, or a group average decrease of >75%.

### Adult *Brugia pahangi* fecundity analysis

*B. pahangi* females lay a mix of embryos, pretzel-stage mf, and mature mf *in vitro*. We designed a computer imaging algorithm to count images of progeny in a high-throughput manner. First, progeny from images from 250 wells were manually counted. Next, we used Fiji^58^ to segment worms using a Sobel filter, Gaussian blur (*σ* = 2.5), and the default iterative selection method for thresholding, and segmented pixels were counted. We split these data into truth and training datasets and used ten-fold cross validation to fit and evaluate a linear model, which explained 88% of the variation in pixel coverage (Supplementary Fig. 6). This model was used to analyze all subsequent wells. Each fecundity value from 48 hpt was normalized to the pretreatment time point to generate a measure of relative fecundity. Parasites that released less than 200 progeny during the overnight recovery period were removed from further analysis. A hit was defined as any compound that caused a decrease of >50% in fecundity in more than two worms, or a group average decrease of >50%.

### Adult *Brugia pahangi* viability analyses

Adult parasites picked into PBS in 96-well plates were stained with CellTox (Promega, Madison, WI) using the manufacturer’s protocol, except using 1/2 the recommended concentration of reagent. After washing twice with PBS, wells were filled to the top with PBS and green fluorescence was imaged at 2X with an ImageXpress Nano. A 14-slice Z-stack was imaged with 250 μm separating each slice, and a maximum projection of the stack was used to assess fluorescence. Worms were segmented with a Sobel filter and Gaussian blur (*σ* = 10), thresholded as above, eroded (x3), and the area of particles with ellipsity of 0-0.5 was measured.

### Adult *Brugia pahangi* metabolic analysis

Conditioned media was thawed and filtered through a 0.2 µm filter plate (Pall, Port Washington, NY) by centrifuging at 1,500 *xg* for 2 min, and filtered media was diluted in PBS to a final dilution of 1:50 for males and 1:150 for females. Lactate production was quantified using the Lactate-Glo kit (Promega), following the manufacturer’s protocol. Luminescence was measured with a SpectraMax i3x plate reader (Molecular Devices). A hit was defined as any compound that caused at least 50% decrease in luminescence in more than two worms, or a group average decrease of >25%.

### Comparative genomics and phylogenetics

Parasite homologs of the human targets of hits in the primary screen were identified with a comparative genomics pipeline. The list of human targets was first expanded using a BLASTp search against the human predicted proteome and retaining hits with percent identity >30%, E-value < 10^−4^, and percent coverage >40%. This expanded list was used for a BLASTp against the predicted proteomes of *B. malayi, Ancylostoma caninum, Ascaris suum, C. elegans, Haemonchus contortus, Strongyloides ratti*, and *Trichuris muris* (all predicted proteomes were downloaded from WormBase ParaSite v16^59^). Hits were filtered as above. Surviving hits were used in a reciprocal BLASTp against the human predicted proteome. Parasite homologs that had top hits to the original human targets were maintained. Target homologs were aligned, trimmed, and phylogenetic trees were inferred with IQ-TREE^60^. Trees were visualized and annotated with ggtree^61^. The entire pipeline can be found at https://github.com/zamanianlab/Phylogenetics/tree/master/Tocris and trees are in Supplementary Data 1.

### Virtual screening against human and parasite AlphaFold2 protein structures

A blind docking procedure was used to predict compound-target binding poses and interaction scores for compounds in the Tocriscreen 2.0 Mini chemical library with AlphaFold2 protein structure models for human and available *B. malayi* orthologs. Scoring features generated from docked compound-target pairs provided input representations for machine learning classifiers. Scripts containing the full comands and all analytical code can be found at https://github.com/zamanianlab/MultivariateScreening-ms.

The compound library was obtained from Tocris in SMILES format along with catalog IDs and UniProt annotations of the known human protein targets. 1,280 SMILES were provided with unique catalog IDs and were sanitized with RDKit (RDKit: Open-source cheminformatics. https://www.rdkit.org) to produce 1,278 RDKit-canonical, isomeric SMILES (Supplementary Table 3). Protonation states were set using *fixpka* utility from OpenEye Scientific, Inc. (QUACPAC 2.2.0.1, Santa Fe, NM). A single 3D conformation was then prepared for each molecule using OpenEye’s *omega2* utility followed by partial charge assignment using *molcharge* with the MMFF method (OMEGA 4.1.1.1: OpenEye Scientific Software, Santa Fe, NM, USA. http://www.eyesopen.com, 2019)^62^. For structures that failed in processing by omega2, the *oeomega* utility was applied using the “*macrocycle*” mode to build conformers. From the 1278 unique RDKit SMILES, a total of 1,534 unique isomers were generated as docking-ready structures in mol2 format. For the Tocris molecules (catalog IDs) having multiple isomers, only the top scoring isomer based on GNINA’s CNNscore was used in further evaluations.

AlphaFold2 model structures were downloaded from: https://alphafold.ebi.ac.uk/downloadfor both human (UP000005640_9606_HUMAN_v2) and *B. malayi* (UP000006672_6279_BRUMA_v2) proteomes. From the human genome, a subset of 3,448 unique Ensembl gene IDs were selected to cover the main drug targets associated with the Tocriscreen library, and the comparative *B. malayi* dataset was generated by retrieving the orthologs of the human genes using biomaRt and the gene trees hosted by WormBas ParaSite^59,63^. To assign target classes, an all vs all BLAST was performed on the human dataset, and a Euclidean distance matrix was calculated using the pairwise bitscores. The targets were then clustered and each cluster was manually annotated to create a total of 114 unique classes. Classes were assigned to *B. malayi* targets based on the known homologies. Using the human medoid member of each class as a reference, target cohorts were structurally aligned using PyMOL’s cealign utility^64^. *B. malayi* orthologs were also aligned onto the human medoids in each target class. Aligned models were then processed using Chimera’s (version 1.16) DockPrep utility and exported in mol2 format^65^.

GNINA was applied in a blind docking strategy, obviating manual binding site specification^66^. For blind docking, more extensive sampling is required (exhaustiveness = 64, rather than default value of 8). The target protein structure was specified as the argument for the *autobox_ligand* option to enable blind docking. CNN scoring was used only for re-scoring final poses by setting *cnn_scoring* option to “rescore.” For CNN-based pose scoring, the “*crossdock_default2018*” model was selected. The top scoring pose based on CNNscore was selected to represent each compound-target pair. All docking runs were performed at the CHTC, which manages job access to Open Science Grid compute resources^67^. Pose locations were calculated as geometric centroid points and stored for docking site analysis on each target class. Points in each target class were clustered by DBSCAN^68^ with ε = 1.5 Å and the *min_samples* threshold scaled for each target class as sqrt(N)/2, where N is the total number of poses (points). Based on pose cluster assignments, an additional scoring feature, site consensus score, was computed for each pose as the fractional occupancy of its pose cluster with respect to all poses observed in target class. For example, if 10% of all poses in a given target class were observed in a given pose cluster, poses in this cluster were assigned a site consensus score of 0.10. Outliers/singleton poses were assigned a site consensus score of 0.0.

Random forest and XGBoost^69^ machine learning models were tuned and fit to the human target docking data using true compound-target pairings. Parameters were tuned using ten-fold cross-validation and evaluated on hold-out test data. The data set (∼1,800 actives and >3.5m decoys) was balanced by downsampling. Models were evaluated using a variety of metrics, including the enrichment factor at 1% (EF1), BEDROC^19^, and RIE^18^, which were implemented into the tidymodels framework using a custom package (https://github.com/zamanianlab/ZamanianLabVSTools)^70^.

### Statistical analyses and replicates

For high-throughput screens, hits were defined as those compounds that had a Z-score > 1 (*Z* = (*x* − µ) ÷ σ). The Z’-factor^13^ of the plate was calculated using the equation: 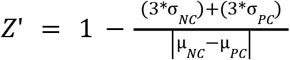. Correlation between raw and subsampled mf videos was assessed using Spearman’s correlation coefficient. Screens against *B. malayi* mf and *C. elegans* N2 were completed once. For *B. malayi* mf dose-response experiments, 4 technical replicates (wells) were included for each concentration, and dose-response curves were fit to a four-parameter log-logistic curve. For *C. elegans* development assays, Dunn’s test for nonparametric multiple pairwise comparisons was used to compare treatment groups to the negative control. For *B. pahangi* adult multivariate experiments, four worms were used for each sex, all coming from a single batch extracted from jirds that were infected on the same day with the same batch of L3s. Comparisons of fecundity between *Brugia* species was calculated using a Kruskal-Wallis one-way ANOVA. A measure of binding pocket dissimilarity (M _d_) was calculated using the equation: 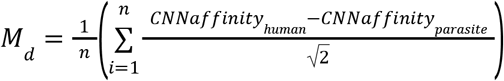, where *CNNaffinity* is a GNINA output and *I* is the compound docked.

## Data and code availability

All analyses and figures can be reproduced using the data (as RDS files) and code available at https://github.com/zamanianlab/MultivariateScreening-ms. All raw imaging data (>2 TB) is available upon request. Docking results and tuned machine learning models are available in a Zenodo repository (https://doi.org/10.5281/zenodo.6599609).

## Acknowledgments

The authors would like to thank Kathy Vaccaro and Elena Rehborg for assistance with *C. elegans* strain maintenance and phenotyping, as well as Gene Ananiev and staff at the UWCCC Small Molecule Screening Facility for assistance with compound library storage and preparation. Parasite materials were provided by the NIH/NIAID Filariasis Research Reagent Resource Center (www.filariasiscenter.org). This research was performed using the compute resources and assistance of the UW-Madison Center For High Throughput Computing (CHTC) in the Department of Computer Sciences. The CHTC is supported by UW-Madison, the Advanced Computing Initiative, the Wisconsin Alumni Research Foundation, the Wisconsin Institutes for Discovery, and the National Science Foundation, and is an active member of the OSG Consortium, which is supported by the National Science Foundation and the U.S. Department of Energy’s Office of Science.

## Supplementary information

**Supplementary Data 1** - Phylogenetic trees that include the human target for all hit compounds along with homologs from selected nematode species

**Supplementary Figure 1** - Assessment of microfilaria congregation in response to variations in handling and environment

**Supplementary Figure 2** - BLAST comparisons of the targets of hit and non-hit compounds

**Supplementary Figure 3** - Dose-responses and pharmacodynamics of hit compounds

**Supplementary Figure 4** - Phylogenetic trees used to select the *C. elegans* orthologs of 6 putative targets

**Supplementary Figure 5** - Batch variability of *Brugia* spp. Fecundity

**Supplementary Figure 6** - Evaluation of a linear model for estimating the number of microfilaria in a well

**Supplementary Table 1** - Comparison of *Brugia malayi* adult and microfilaria RNA-seq profiles

**Supplementary Table 2** - List of the *B. malayi* target structure that were estimated by docking to be the most likely target for hit compounds

**Supplementary Table 3** - Tocriscreen 2.0 compounds in sanitized SMILES format used for molecular docking

