## Supplementary figures and images for "Multivariate chemogenomic screening prioritizes new macrofilaricidal leads"

### Supplementary Fig 1

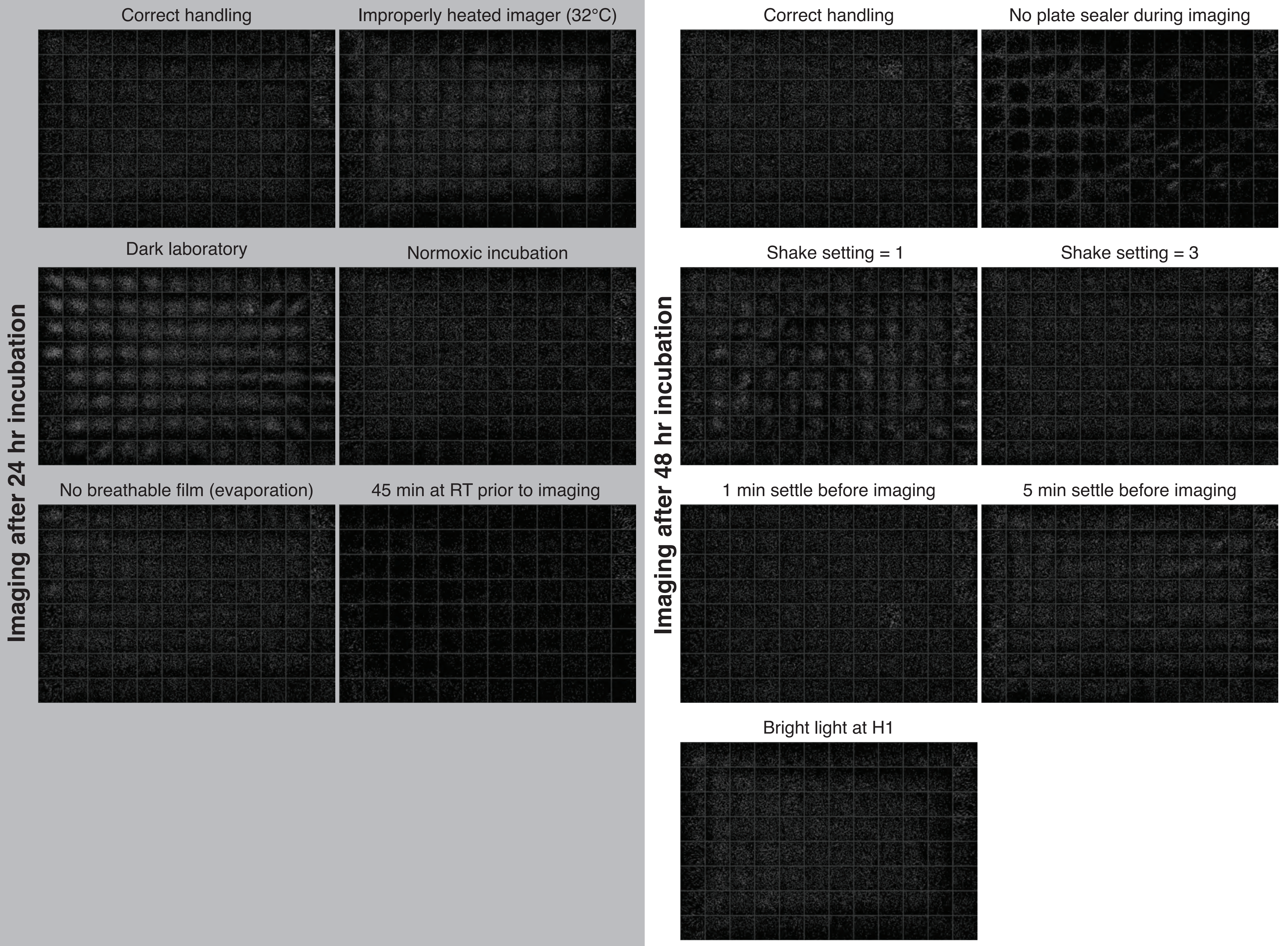

### Supplementary Fig 2

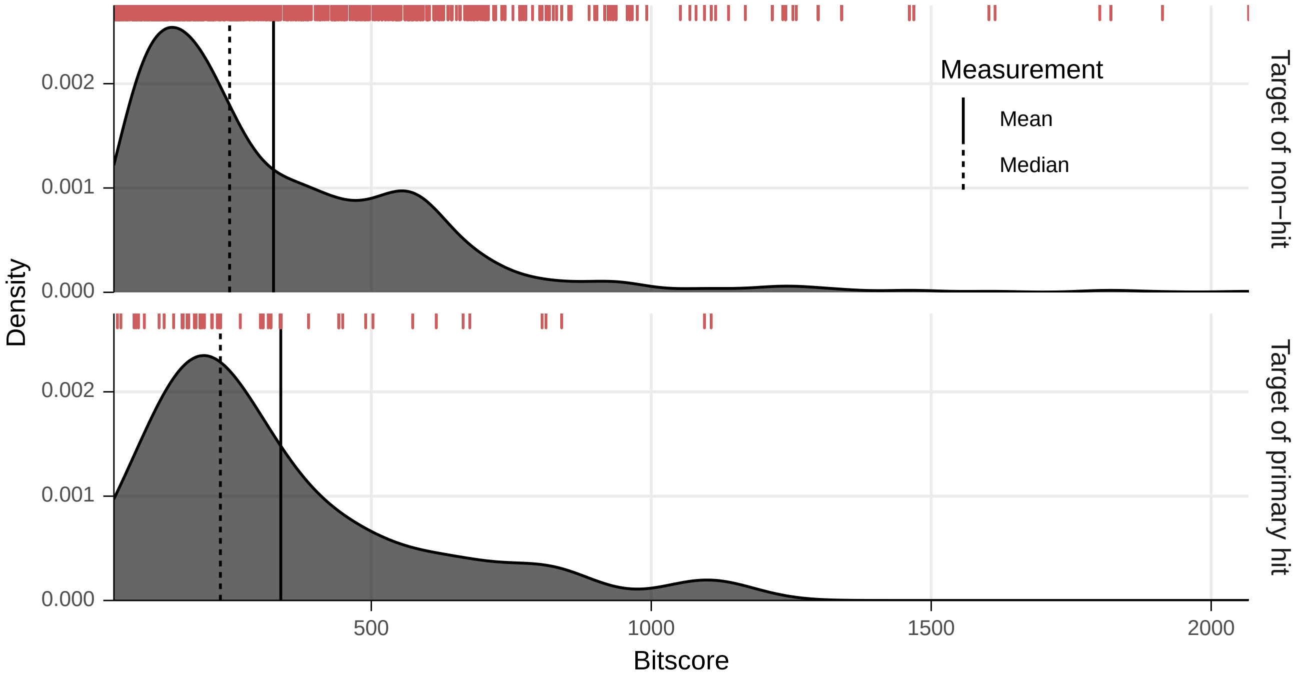

### Supplementary Fig 3

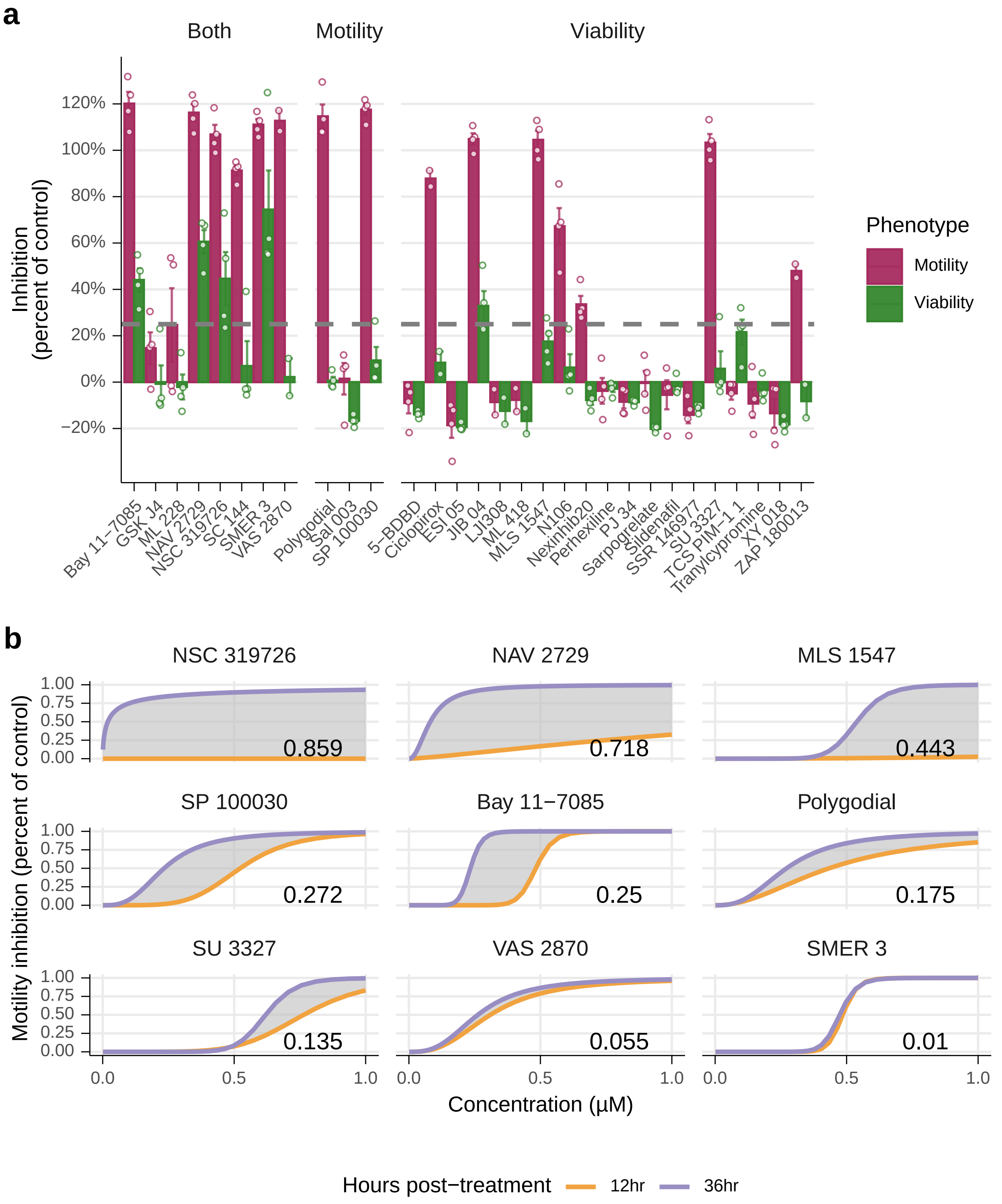

### Supplementary Fig 4

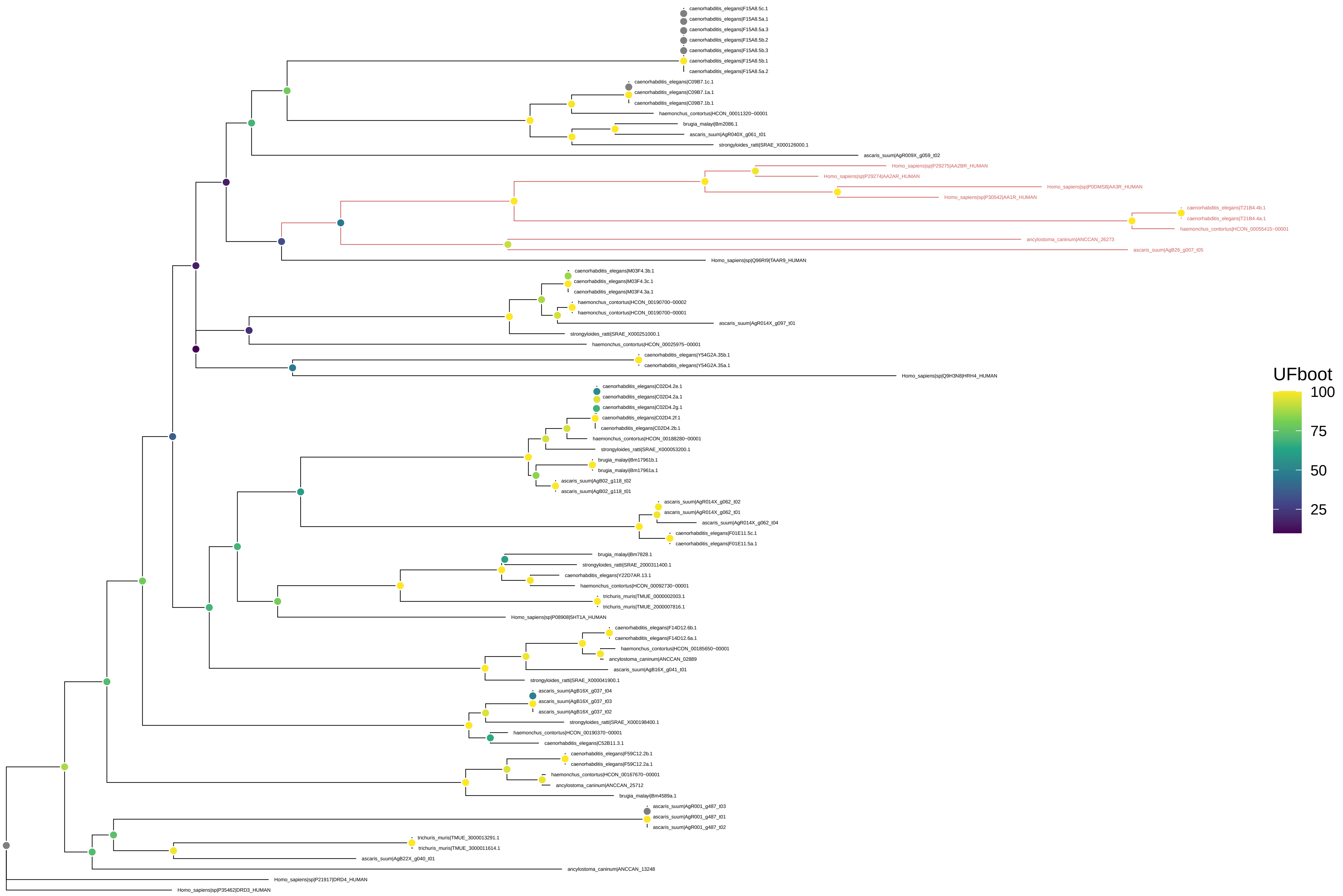

ARF6

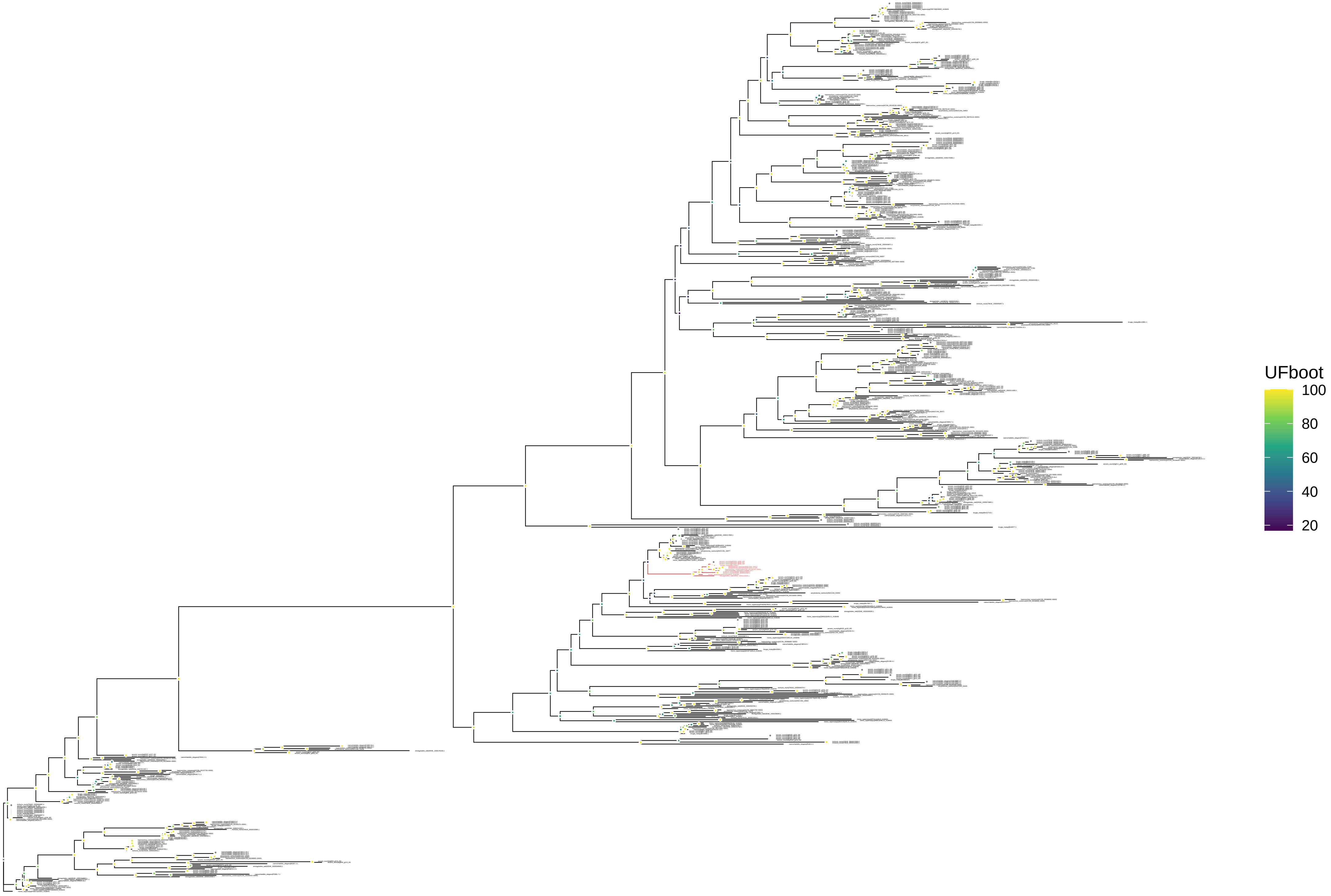

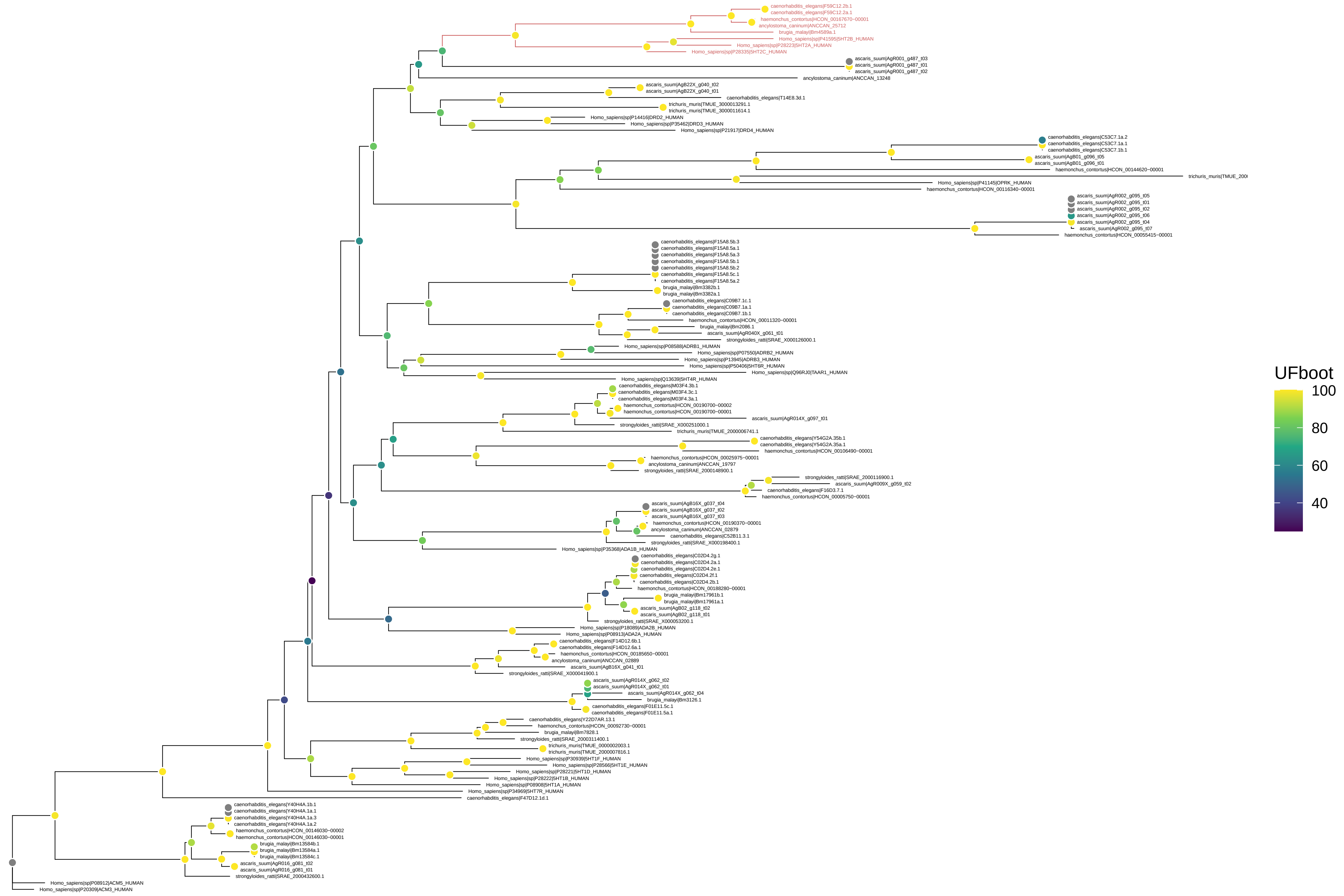

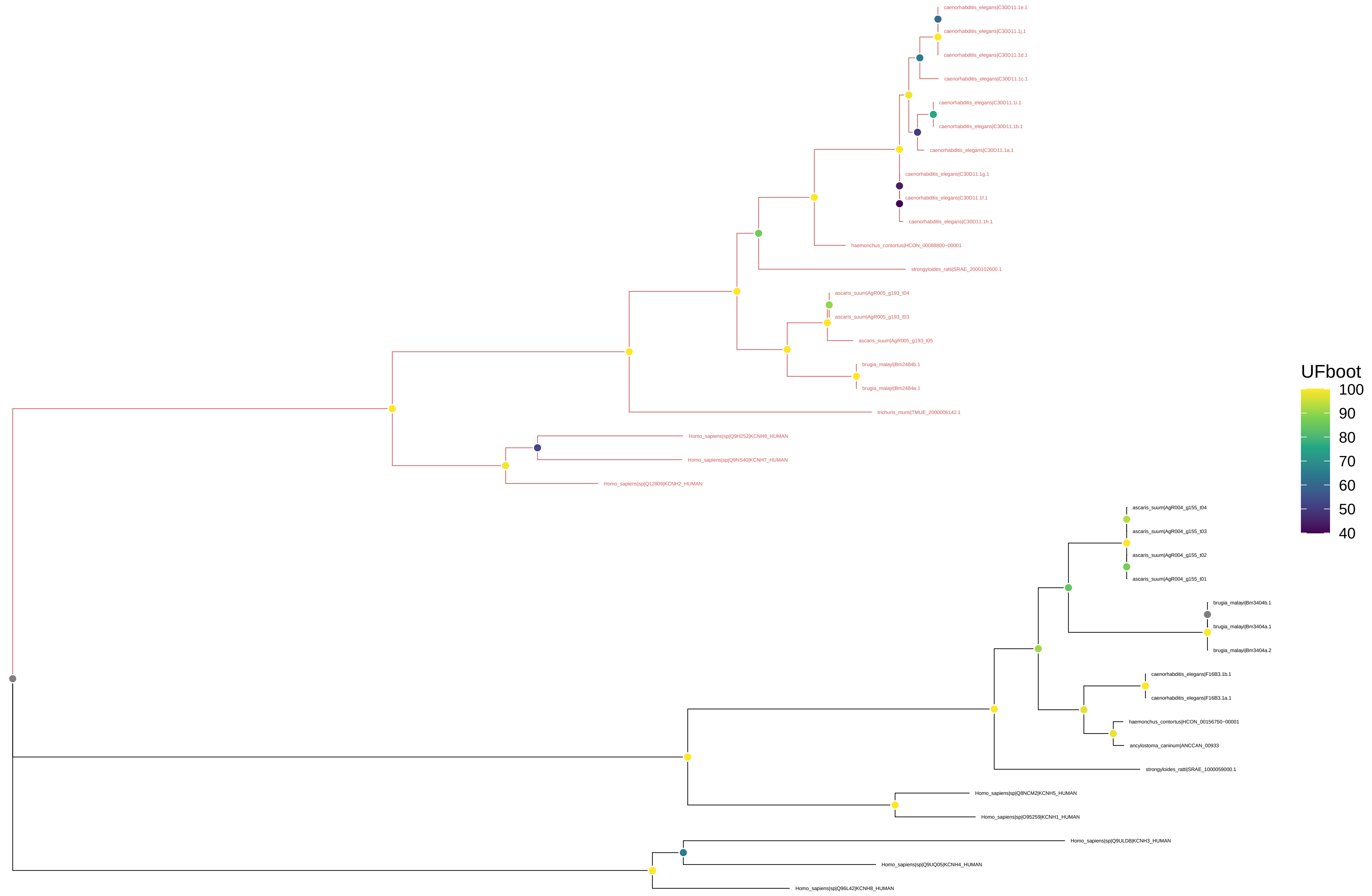

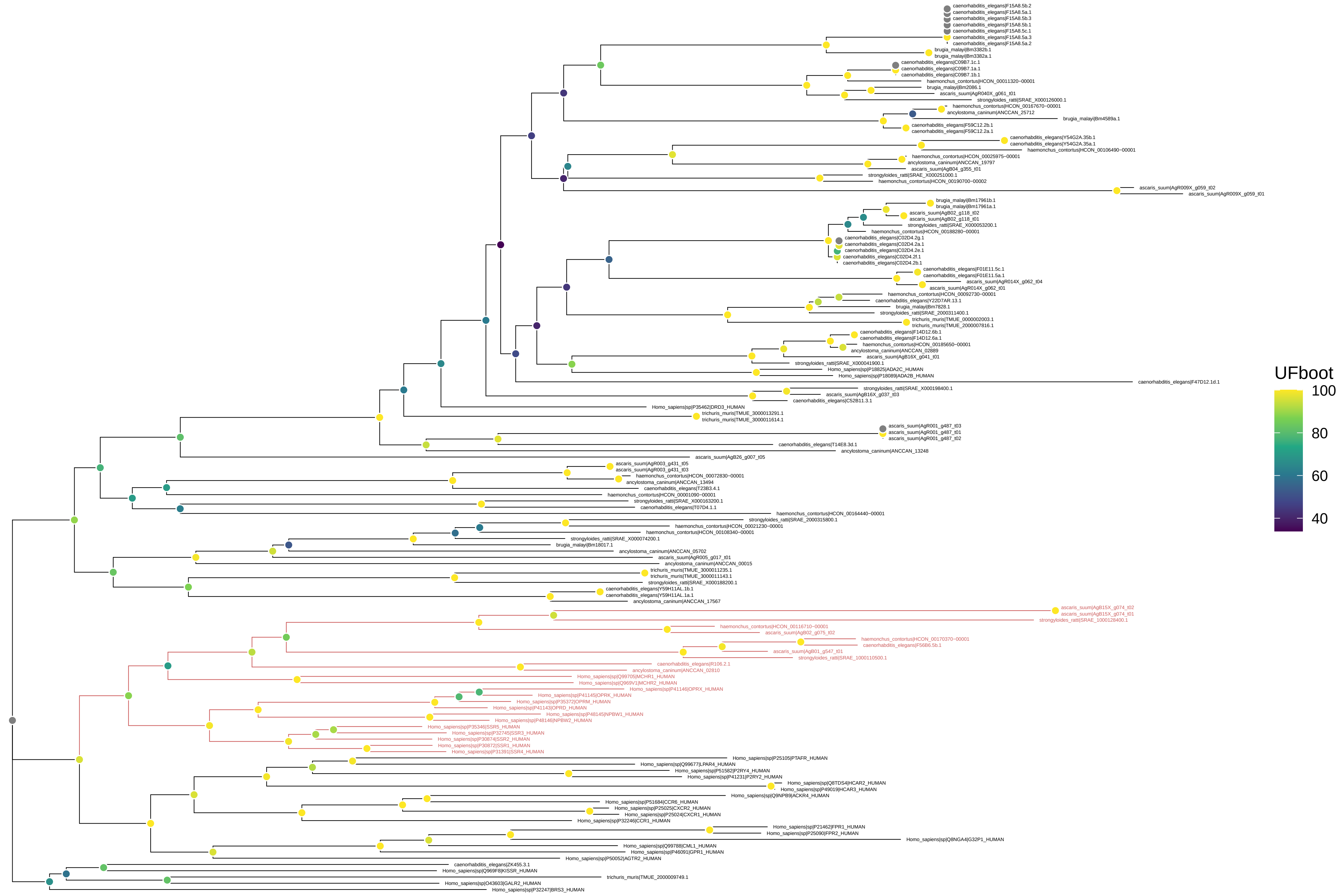

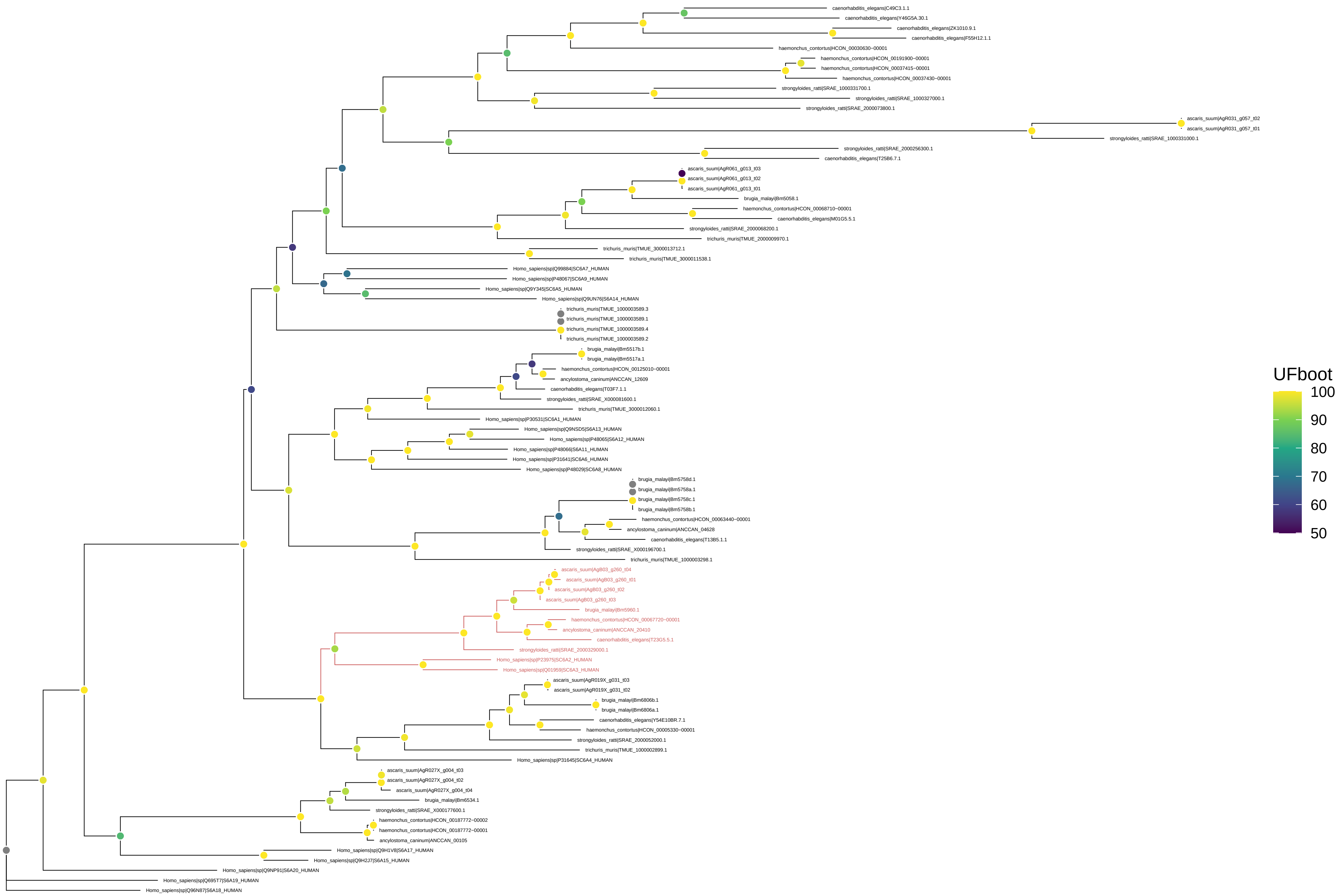

### Supplementary Fig 5

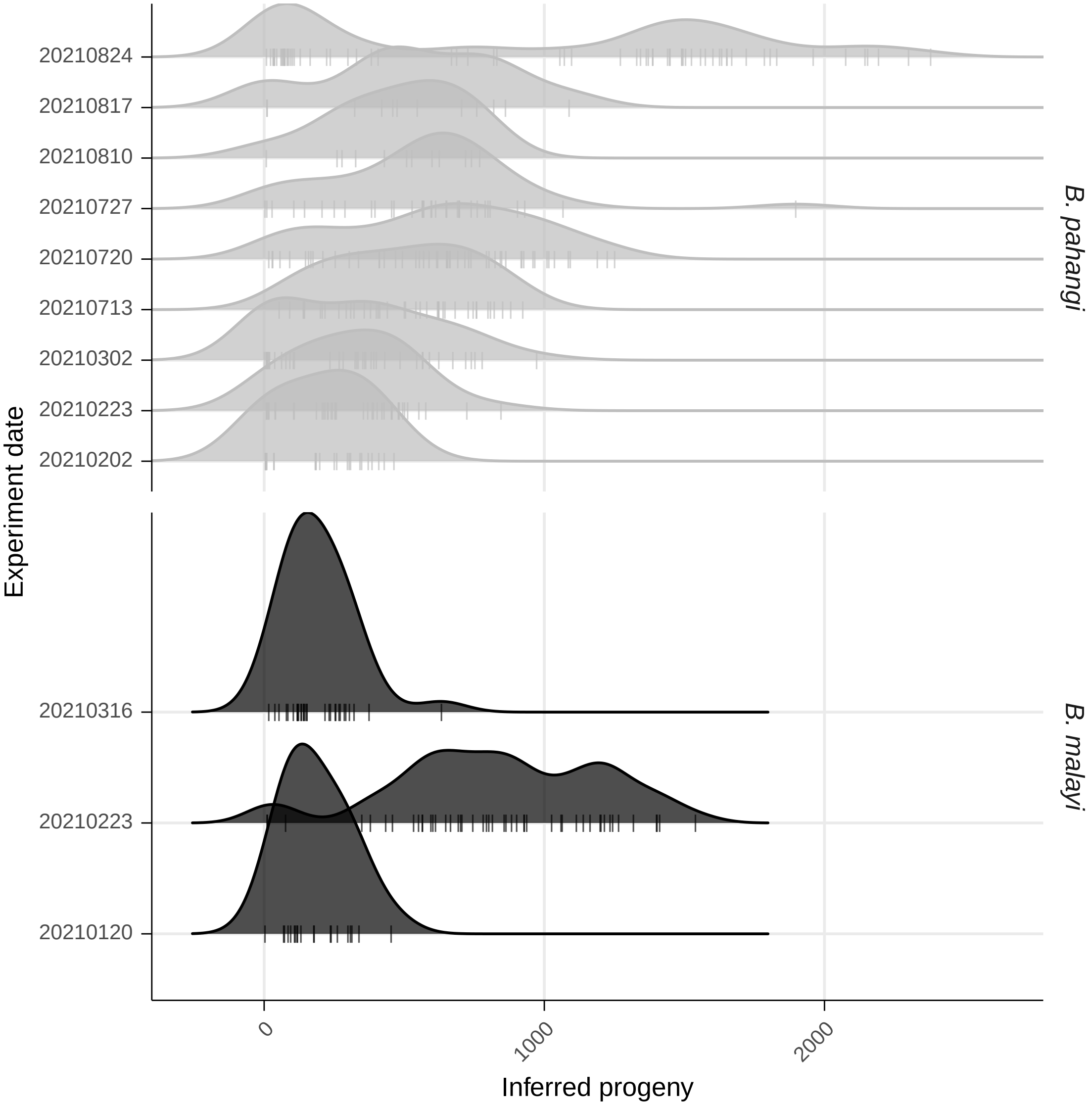

### Supplementary Fig 6

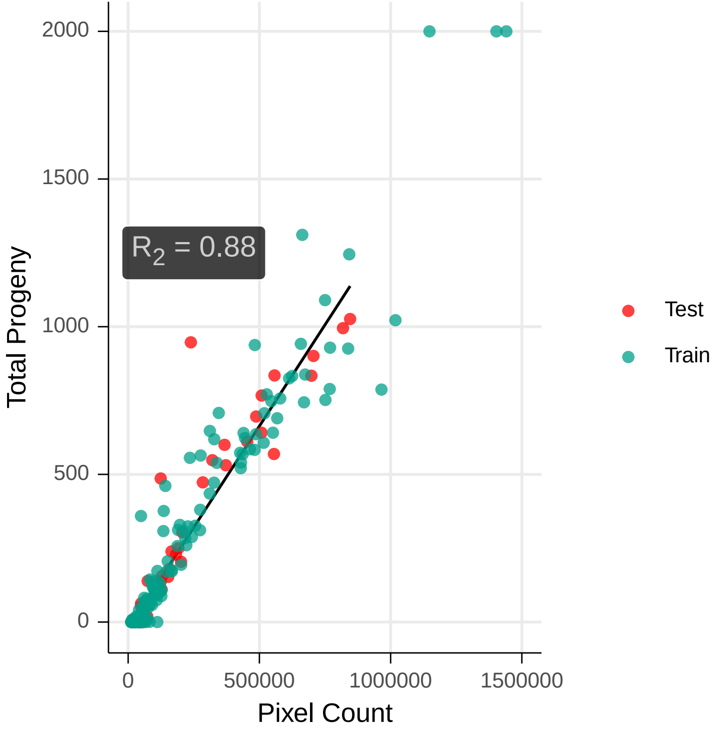
